# Sialin2 Functions as a Mammalian Nitrate Sensor to Sustain Mitochondrial Homeostasis

**DOI:** 10.1101/2025.05.04.652104

**Authors:** Xiaoyu Li, Songyue Wu, Zichen Cao, Ou Jiang, Shaorong Li, Xinyue Chen, Yao Feng, Bo Zhou, Chen Zhang, Guozhi Xiao, Jinsong Wang, Jian Zhou, Mo Chen, Renhong Yan, Songlin Wang

**Affiliations:** Salivary Gland Disease Center and Beijing Key Laboratory of Tooth Regeneration and Function Reconstruction, Beijing Laboratory of Oral Health and Beijing Stomatological Hospital, Capital Medical University, Beijing 100069, China; Department of Biochemistry and Molecular Biology, Capital Medical University School of Basic Medicine, Beijing 100069, China; Department of Biochemistry, Key University Laboratory of Metabolism and Health of Guangdong, SUSTech Homeostatic Medicine Institute, School of Medicine, Southern University of Science and Technology, Shenzhen 518055, China; Department of Pharmacology, Joint Laboratory of Guangdong-Hong Kong Universities for Vascular Homeostasis and Diseases, SUSTech Homeostatic Medicine Institute, School of Medicine, Southern University of Science and Technology, Shenzhen 518055, China; Department of Biochemistry, SUSTech Homeostatic Medicine Institute, School of Medicine, Shenzhen Key Laboratory of Cell Microenvironment, Guangdong Provincial Key Laboratory of Cell Microenvironment and Disease Research, Southern University of Science and Technology, Shenzhen, 518055, China; Department of Endodontics, Beijing Stomatological Hospital, Capital Medical University, Beijing 100071, China

**Keywords:** Nitrate, Sialin2, sensor, AMPK, mitochondria

## Abstract

Nitrogen homeostasis is fundamental for cellular physiology, yet mammalian nitrate (NO3^−^) sensing mechanisms remain elusive. Here, we identify Sialin2—a proteolytic fragment of the nitrate transporter Sialin generated by cathepsin B (CTSB) cleavage— as the first mammalian nitrate sensor. Microscale thermophoresis (MST) reveals Sialin2 as a high-affinity nitrate sensor, while Cryo-Electron Microscopy (Cryo-EM) uncovers its structural basis for signaling. We show that Sialin2 localizes to mitochondria and scaffolds liver kinase B1 (LKB1)-AMP-activated protein kinase (AMPK) complexes to drive organelle-specific metabolic adaptation via spatiotemporally controlled AMPK activation, enhancing mitochondrial biogenesis, ATP production, and cell survival. Real-time tracking using the sCiSiNiS biosensor demonstrates nitrate signaling dynamics at physiological levels. This signaling axis redefines nitrate as a direct ligand activating receptor-like cascades, independent of classical nitric oxide synthesis. Our findings establish a paradigm of “inorganic salt signaling biology”, wherein anions co-opt trafficking systems to achieve signaling specificity, offering therapeutic avenues for mitochondrial disorders.

## Introduction

Nitrogen is an essential element for all living organisms and a fundamental component of numerous cellular molecules, including nucleic acids, amino acids, proteins, and phospholipids^1^. Given its fundamental roles in cellular physiology, organisms must precisely regulate nitrate homeostasis to sustain essential biological processes^2^. Among nitrogen species, nitrate (NO3^−^) has emerged as a biologically active molecule with dual roles as both a nutrient and a signaling agent^3^. In recent decades, nitrate has been implicated in a wide range of physiological processes, including enhancing gastrointestinal blood flow^4,5^, modulating fat metabolism to reduce obesity^6^, increasing the sensitivity of cisplatin chemotherapy in tumor therapy^7^, and mitigating radiation-induced damage to salivary glands^8–10^. These diverse roles underscore the importance of nitrate in maintaining organismal health and homeostasis.

Despite its significance, the mechanisms by which mammalian cells sense and respond to nitrate remain poorly understood. In bacteria and plants, nitrate sensing is well-documented, with specialized receptors and signaling pathways that precisely control nitrogen metabolism and adaptive responses under nutrient or stress conditions^11–13^. In contrast, mammalian nitrate perception has not been delineated, limiting our understanding of its role in cellular signaling networks. Our recent study has identified that Sialin, a mammalian nitrate transporter encoded by the *SLC17A5* gene, is responsible for nitrate uptake and transport^14^. Primarily localized to the plasma and lysosomal membranes, Sialin facilitates nitrate homeostasis and has been implicated in various physiological and pathological processes, including lysosomal function regulation, metabolic adaptation under stress conditions, and neurodegeneration diseases^14–17^. However, unresolved questions remain about whether Sialin or other proteins function as nitrate sensors and how nitrate signaling is integrated into cellular homeostasis. Addressing this knowledge gap is critical for advancing our understanding of nitrate-mediated biological processes and their potential therapeutic applications.

In this study, we address this gap by revealing that nitrate exposure triggers proteolytic cleavage of Sialin by cathepsin B (CTSB), generating a distinct fragment termed Sialin2, which functions as the first identified mammalian nitrate sensor. We further demonstrate that Sialin2 localizes to mitochondria, where it initiates a spatially compartmentalized signaling cascade involving liver kinase B1 (LKB1) and AMP-activated protein kinase (AMPK), directly linking nitrate sensing to mitochondrial biogenesis, metabolic adaptation, and cellular survival. Our findings thus unveil a previously unrecognized mode of inorganic nitrate signaling, establishing Sialin2 as a cellular homeostasis regulator in nutrient signaling networks and providing a mechanistic foundation for therapeutic exploration targeting metabolic and degenerative diseases.

## Results

### Nitrate-induced CTSB-Mediated Cleavage of Sialin Generates Mitochondria-Localized Sialin2

To elucidate the molecular mechanisms of nitrate sensing and signaling, we sought to identify key proteins involved in nitrate detection and subsequent signal transduction. Immunoblot analysis of tissues from Alzheimer’s disease (AD), radiation-damaged, high-fat diet (HFD)-induced obesity (OB), and estrogen-deficiency-induced osteoporosis (OP) mouse models^8,18–20^ revealed a consistent increase in a ∼31 kDa protein fragment, designated Sialin2, derived from full-length Sialin (∼54 kDa), in the cerebral cortex, salivary glands, white fat, and femur (Fig. 1a, Extended Data Fig. S1a). Notably, *SLC17A5* mRNA level remained unchanged across all conditions, ruling out transcriptional regulation as a contributing factor (Extended Data Fig. S1b). Immunoprecipitation in HEK293T cells confirmed the endogenous presence of Sialin2 (Fig. 1b), and *Slc17a5* knockout (KO) mice exhibited a complete loss of both Sialin and Sialin2, confirming antibody specificity and the origin of Sialin2 (Extended Data Fig. S1c).

**Figure 1.**
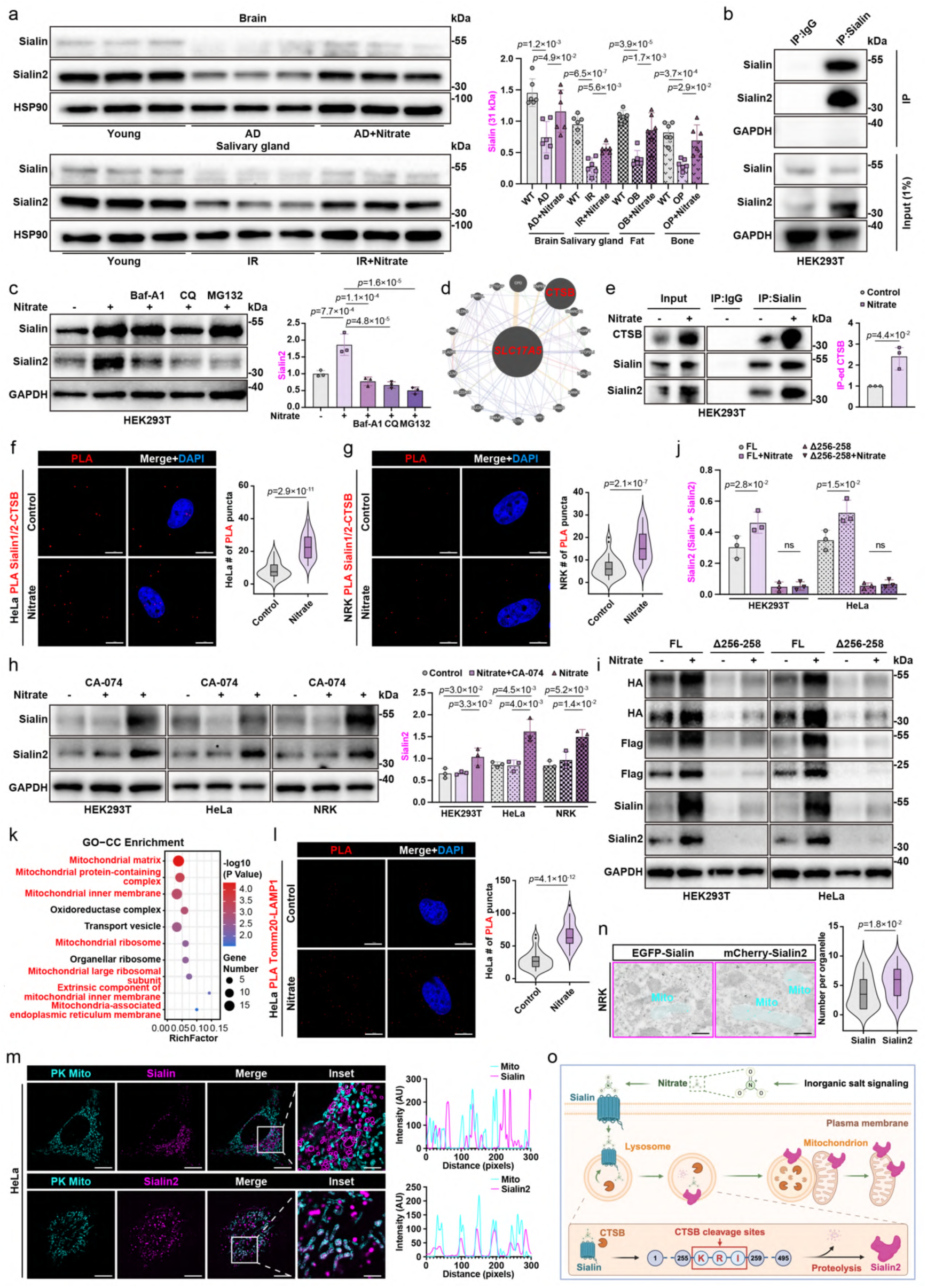
Nitrate-induced CTSB-dependent cleavage of Sialin at K256/R257/I258 generates Sialin2, which localizes to mitochondria a, Immunoblot analysis of Sialin and Sialin2 protein normalized to HSP90 in cerebral cortex from Alzheimer’s disease (AD) mice, salivary glands from irradiation (IR)- damaged mice. Representative data of n = 6 independent experiments were shown. See full immunoblot images and quantitation in Extended Data Fig. 1a. b, Endogenous Sialin2 was immunoprecipitated (IP’ed) from HEK293T cells, with IgG as negative control. Representative data of n = 3 independent experiments were shown. c, Immunoblot analysis of Sialin and Sialin2 in HEK293T cells pre-treated for 6 h with DMSO (vehicle), 200 nM Baf-A1/40 μM CQ (lysosomal inhibitors), or 10 μM MG132 (proteasome inhibitor), followed by 4 h stimulation with 4 mM nitrate. Representative images of n = 3 independent experiments were shown. d, Protein-protein interaction network (GeneMANIA) predicted cathepsin B (CTSB) as a top candidate protease interacting with Sialin. e, Co-IP of Sialin and CTSB in HEK293T cells treated with 4 mM nitrate for 4 h. Representative data of n = 3 independent experiments were shown. f, g, Proximity ligation assay (PLA) of Sialin/Sialin2-CTSB in HeLa (f) or NRK (g) cells treated with nitrate (4 mM, 4 h). N = 30 cells from representative experiments of three repeats. h, Immunoblot analysis of Sialin and Sialin2 in HEK293T (left panel), HeLa (middle panel) or NRK (right panel) cells pre-treated for 6 h with a vehicle control (DMSO) and 20 μM CTSB inhibitor CA-074, followed by 4 h stimulation with 4 mM nitrate. Representative images of n = 3 independent experiments were shown. i, j, HEK293T (left panel) or HeLa cells (right panel) introduced with HA/Flag-tagged full length (FL) Sialin or mutant full-length Sialin (KRI/AAA) were treated with nitrate (4 mM, 4 h), followed by analysis of HA, Flag, Sialin, and Sialin2. Representative images of n = 3 independent experiments were shown. I, Ile; K, Lys; R, Arg. k, Top enriched cellular component GO terms for unique Sialin2-related proteomes in HEK293T cells listed by the rank of *P* values based on DAVID analysis. l, PLA of Tomm20-LAMP1 in HeLa cells treated with nitrate (4 mM, 4 h). N = 30 cells from representative experiments of three repeats. m, High-sensitivity structured illumination microscopy (HiS-SIM) live-cell images of Sialin or Sialin2 colocalized with MitoTracker-labeled mitochondria (PK Mito) in HeLa cells stably expressed GFP-Sialin or GFP-Sialin2. See quantification in Extended Data Fig. 2f. n, Immunoelectron microscopy images of Sialin or Sialin2 localized to mitochondria in NRK cells stably expressed GFP-Sialin and mCherry-Sialin2. N = 30 cells from representative experiments of three repeats. o, Schematic illustration of nitrate-activated CTSB cleaving Sialin at K256/R257/I258 to generate Sialin2, which is preferentially localized to mitochondria. For all panels, data are represented as mean ± SD, *P* value denotes *t*-test. Scale bar: 10 μm.

In response to nitrate treatment (4 mM, 4 h), Sialin2 levels were substantially increased in HEK293T, HeLa, and NRK cells. However, inhibiting lysosomal degradation (Baf-A1/CQ) or proteasomal degradation (MG132)^21,22^ markedly reduced Sialin2 formation, suggesting distinct contributions of these pathways (Fig. 1c, Extended Data Fig. S1d-f). To systematically identify the protease responsible for Sialin cleavage, we performed an in-silico protease substrate prediction and protein-protein interaction network analysis (GeneMANIA), which prioritized cathepsin B (CTSB) as the top candidate interacting with Sialin (Fig. 1d). Co-immunoprecipitation (Co-IP) assays confirmed a direct interaction between CTSB and Sialin in nitrate-treated HEK293T cells (Fig. 1e). Proximity ligation assays (PLA) further demonstrated enhanced interaction between Sialin and CTSB in nitrate-treated HeLa and NRK cells, with significantly increased signal intensity compared to untreated controls (Fig. 1f, g). Critically, pretreatment with the CTSB-specific inhibitor CA-074 completely blocked nitrate-induced Sialin2 generation in all tested cell lines (Fig. 1h, Extended Data Fig. S1d) without altering *SLC17A5* transcript levels (Extended Data Fig. S1e, f).

To define the molecular determinants of CTSB-mediated Sialin processing, we performed site-directed mutagenesis combined with protease substrate prediction^23^. Sequence analysis of Sialin identified a conserved K256/R257/I258 (KRI) motif near the cleavage site that produces Sialin2 (Extended Data Fig. S1h), aligning with CTSB’s canonical cleavage preference. We expressed HA/Flag-tagged full-length Sialin or a KRI/AAA mutant, site-directed mutagenesis of residues Lys^2^^56^, Arg^2^^57^, and Ile^258^ (K256, R257, I258) to alanine (Extended Data Fig. S1g), in HEK293T and HeLa cells. Nitrate treatment induced robust cleavage of full-length Sialin, generating Sialin2 (Fig. 1i, j). In contrast, the KRI/AAA mutant exhibited complete resistance to CTSB-mediated proteolysis. Although basal expression of the mutant was moderately reduced compared to wild-type Sialin, cleavage efficiency was normalized to total Sialin levels (Sialin2/[Sialin+Sialin2]), and residual mutant protein remained detectable (Fig. 1 i, j). Direct expression of the predicted cleavage product (recombinant Sialin2-Flag), yielded a stable protein with a molecular weight consistent with endogenous Sialin2, confirming that the KRI-proximal sequence encodes the authentic cleavage product (Extended Data Fig. S1i, j).

Since that Sialin2 is generated through CTSB-mediated proteolysis, we next investigated its functional significance, focusing on its subcellular localization. Gene ontology (GO) analysis of Sialin2-associated proteomes in HEK293T cells revealed significant enrichment for mitochondrial components (Fig. 1k, Extended Data Fig. S2a, b), suggesting a potential role in mitochondrial function. Proximity ligation assays (PLA) demonstrated that nitrate treatment increased mitochondrial-lysosomal contacts (Tomm20-LAMP1 proximity), suggesting a potential translocation mechanism for Sialin2 (Fig. 1l, Extended Data S2c–e)^24,25^. High-sensitivity structured illumination microscopy (HiS-SIM) of live HeLa cells expressing GFP-Sialin2 showed strong colocalization with MitoTracker-labeled mitochondria, whereas full-length Sialin primarily localized to lysosomes (Fig. 1m, Extended Data Fig.S2f, g). Immunofluorescence (IF) and immunoelectron microscopy (IEM) in NRK cells stably expressing GFP-Sialin and mCherry-Sialin2 confirmed that Sialin2, but not full-length Sialin, preferentially localized to mitochondria, while also partially distributed in lysosomes, endoplasmic reticulum, and Golgi apparatus (Fig. 1n, Extended Data S3a– i).

Collectively, these findings demonstrate that nitrate activates CTSB, which cleaves Sialin at the K256/R257/I258 (KRI) motif, generating Sialin2, which is targeted to mitochondria (Fig. 1o). The conservation of this cleavage event across diverse pathophysiological conditions, including neurodegenerative disease, metabolic disorders, and tissue damage, suggests a universal role for Sialin2 in nitrate sensing and cellular adaptation^24,25^. The mitochondrial localization of Sialin2 positions it as a spatial regulator that links extracellular nitrate levels to organelle-specific responses, offering new insights into nitrate-dependent metabolic regulation and cellular homeostasis.

### Structural and Functional Characterization of Sialin2 as a Nitrate Sensor

To elucidate the protein conformation and molecular basis of nitrate sensing by Sialin2, we purified recombinant Sialin2 using affinity chromatography and size-exclusion chromatography (SEC), yielding a monodisperse peak indicative of a homogeneous preparation (Fig. 2a–c). The Sialin2 protein was analyzed by cryo-electron microscopy (cryo-EM), and an attempt was made to elucidate its structure (Fig. 2f, Extended Data Fig. S4a–c)^26^, representative cryo-EM micrographs and 2D class averages displayed well-defined particles and six transmembrane helices. Although the small size of Sialin2 (∼35 kDa) limited the resolution of high-resolution structural details, we successfully refined the density of the six transmembrane helices (Fig. 2d, e). Experimental characterization revealed that Sialin2 forms smaller micelle-embedded particles compared to full-length Sialin. The structural analysis demonstrated significant divergence from the AlphaFold2-predicted conformation, particularly in transmembrane helices TM3, TM4, and TM6, which adopt a more compact structural arrangement than predicted (Fig. 2f, g, Extended Data Fig. S5a–d). These findings provide the first structural insights into Sialin2, highlighting its unique architecture and potential nitrate-binding site.

**Figure 2.**
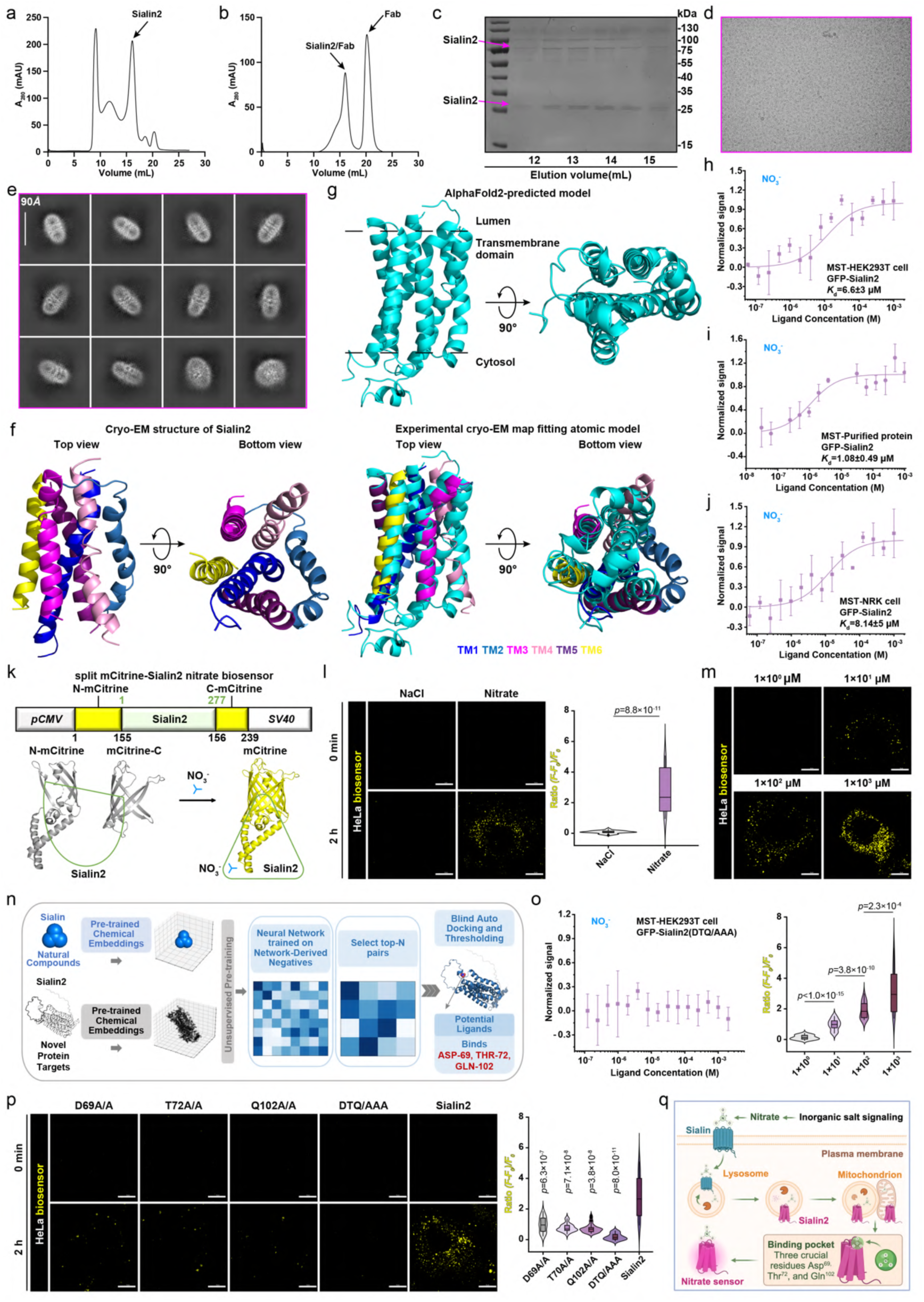
Structural and functional characterization of Sialin2 as a nitrate sensor. a, b, Size exclusion chromatography (SEC) final purification profiles of recombinant Sialin2 (a) and Sialin2/Sialin2-Fab complex (b). c, Coomassie blue-stained SDS–PAGE of purified Sialin2, small multi-transmembrane domain proteins exhibit gradient multimerization patterns (35 kDa, 70 kDa, 105 kDa, etc.) on SDS-PAGE while maintaining homogeneous monomeric states in solution as shown in (d). d, e, Representative cryo-EM micrograph (d) and 2D class averages (e) of Sialin2. f, Cryo-EM fitting-model of Sialin2. g, Experimental cryo-EM map fitting-model of Sialin2 compared to AlphaFold2-predicted model. h–j, Nitrate binding affinity (*K*d) of GFP-Sialin2 measured by microscale thermophoresis (MST) in HEK293T cell lysate (6.6 ± 3 μM; h), purified protein (1.08 ± 0.49 μM; i), and NRK cell lysate (8.14 ± 5 μM; j). N = 3. k, Schematic representation of domain structure and design of the split mCitrine-Sialin2 nitrate biosensor (sCiSiNiS). Nitrate triggered a conformational change that restored mCitrine fluorescence. l, m, Confocal live-cell imaging of cytoplasmic nitrate detected by sCiSiNiS in HeLa cells treated with nitrate (4 mM; l) or dose-dependent nitrate (0–1000 μM; m) for 2 h. Fluorescence intensity normalized as (*F* - *F*0)/*F*0. N = 30 cells from representative experiments of three repeats. n, AI-Bind predicted nitrate-binding sites (red) on Sialin2. o, GFP-Sialin2-DTQ/AAA mutant (D69A/T72A/Q102A) abolished nitrate binding in HEK293T cell lysate. N = 3. D, Asp; T, Thr; Q, Gln. p, Confocal live-cell imaging of sCiSiNiS mutants (D69A, T72A, Q102A, or DTQ/AAA) in HeLa cells treated with nitrate (4 mM) or NaCl for 2 h. Fluorescence intensity normalized as (*F* - *F*0)/*F*0. N = 30 cells from representative experiments of three repeats. q, Schematic illustration of Sialin2 as a nitrate sensor that directly binds nitrate. For all panels, data are represented as mean ± SD, *P* value denotes *t*-test. Scale bar: 10 μm.

To investigate the functional role of Sialin2’s multi-transmembrane architecture, we conducted microscale thermophoresis (MST) experiments. The results demonstrated that GFP-tagged Sialin2 specifically binds nitrate with high affinity in HEK293T cell lysate samples. GFP-tagged Sialin2 specifically bound nitrate with high affinity, demonstrating dissociation constants (*K*d) of 6.6 ± 3 μM in HEK293T cell lysate (Fig. 2h, Extended Data Fig. S6a) and 1.08 ± 0.49 μM in purified recombinant protein samples (Fig. 2i). Comparable affinity was observed in NRK cell lysates (*K*d = 8.14 ± 5 μM; Fig. 2j)^27,28^. In contrast, full-length Sialin displayed significantly lower nitrate-binding affinity (*K*d = 342 ± 209 μM in lysate, *K*d = 7.34 ± 3.84 μM in purified recombinant protein; Extended Data Fig. S6a, h, i), suggesting that proteolytic cleavage either exposes or reconfigures the nitrate-binding site. Specificity assays confirmed that Sialin2 not GFP binds nitrate directly, with no interaction observed for chlorate (ClO3^−^, structurally related anions), sialic acid (SA), asparatic acid (ASP), glutamic acid (Glu), or Na^+^ controls (Extended Data Fig. S6b–g, j, k).

Leveraging these structural and biochemical insights, we engineered a split mCitrine-Sialin2 nitrate biosensor (sCiSiNiS) that reconstitutes fluorescence upon nitrate-induced conformational changes (Fig. 2k)^13,29^. Time-resolved confocal imaging in live HeLa cells revealed a rapid nitrate-specific response, with sCiSiNiS fluorescence intensity increasing within 2 hours of 4 mM nitrate treatment, whereas equimolar NaCl elicited no signal (Fig. 2l). The biosensor exhibited a dynamic range of 0 to 1 mM nitrate (Fig. 2m), consistent with the *in vitro* MST-derived *K*d values. Similar responses were observed in NRK cells (Extended Data Fig. S6l, m), highlighting the broad applicability of this tool.

To define the molecular basis of nitrate recognition, we employed AI-Bind, a deep learning-based predictor^30,31^. Structural analysis identified three crucial residues Asp^69^, Thr^72^, and Gln^102^ (D69, T72, Q102), within the polar binding pocket that are essential for nitrate recognition (Fig. 2n, Extended Data Fig. S7a). Site-directed mutagenesis of these residues to alanine (D69A, T72A, Q102A, or DTQ/AAA; Extended Data Fig. S7b, c) abolished nitrate binding in MST assays (Fig. 2o) and eliminated sCiSiNiS fluorescence activation in HeLa and NRK cells (Fig. 2p, Extended Data Fig. S7e), confirming their essential role in nitrate recognition. Importantly, this functional impairment was independent of protein stability, as immunoblot analysis verified equivalent expression levels of wild-type and mutant Sialin2 (Extended Data Fig. S7d). Collectively, our structural and functional analyses establish Sialin2 as a high-affinity nitrate sensor with a precisely defined molecular architecture. The development of sCiSiNiS enables real-time nitrate detection, and the identification of key nitrate-binding residues provides a mechanistic framework for understanding nitrate signaling (Fig. 2q).

### Nitrate-Sialin2 Signaling Recruits LKB1 to Mitochondria for Compartmentalized AMPK Activation

To identify downstream metabolic targets of nitrate signaling, we performed phosphoproteomic profiling in HEK293T cells treated with 4 mM nitrate (Fig. 3a, Extended Data Fig. S8a). AMPK signaling emerged as the top enriched pathway (Fig. 3b)^33,34^, consistent with a marked increase in AMPK phosphorylation at Thr172 (pAMPK^T172^) in nitrate-treated NRK and HEK293T cells (Fig. 3c, Extended Data Fig. S8b, c). Notably, *SLC17A5* knockout abolished nitrate-induced AMPK activation, a defect fully rescued by wild-type Sialin2 but not the KRI/AAA mutant (Fig. 3c, d, Extended Data Fig. S8d–f). This nitrate-sensing function of Sialin2 is specific and non- redundant, as *SLC17A5* knockout did not impair AMPK activation under glucose starvation or AICAR treatment (Extended Data Fig. S8g).

**Figure 3.**
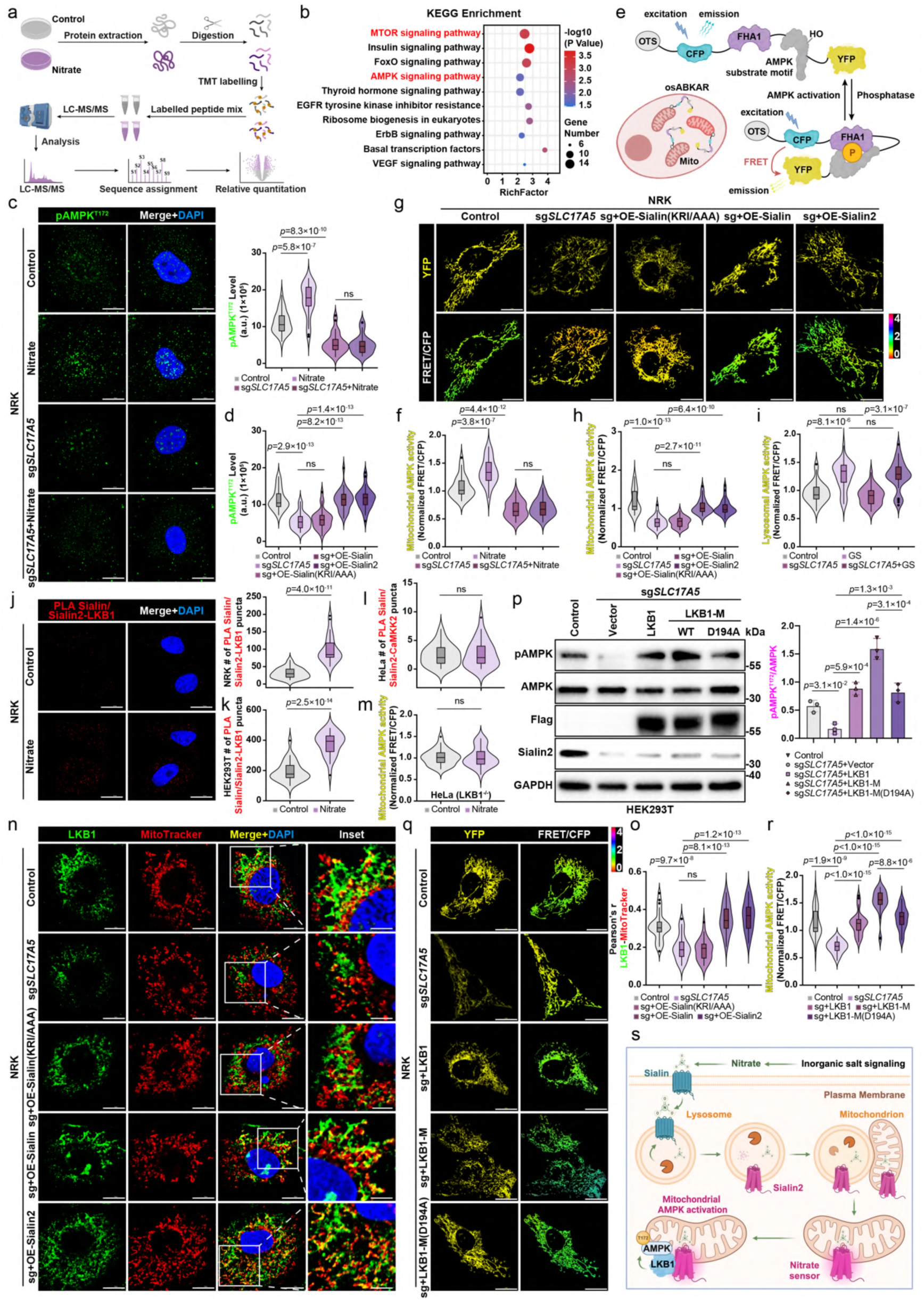
Nitrate-Sialin2 signaling promotes LKB1-mitochondrial recruitment to activate AMPK phosphorylation a, Schematic workflow of phosphoproteomic analysis. b, Top enriched KEGG pathways in HEK293T cells treated with 4 mM nitrate or control vehicle for 4 h listed by the rank of *P* value based on DAVID analysis. Data represent three biological replicates per condition. c, IF staining images (left) and quantification (right) of pAMPK^T172^ in control and *SLC17A5* knockout (sg*SLC17A5*) NRK cells treated with nitrate (4 mM, 4 h). N = 30 cells from representative experiments of three repeats. d, Quantification of pAMPK^T172^ in control and sg*SLC17A5* NRK cells reconstituted with Sialin (KRI/AAA), Sialin, and Sialin2. N = 30 cells from representative experiments of three repeats. See images in Extended Data Fig. 8e. e, Schematic diagram of subcellular compartment-specific biosensor osABKAR. f, Quantification of FRET/CFP ratio of Mito-ABKAR in control and sg*SLC17A5* NRK cells treated with nitrate (4 mM, 4 h). N = 30 cells from representative experiments of three repeats. See images in Extended Data Fig. 9d. g, h, IF staining images (upper, representative YFP images, lower, representative pseudocolor images of FRET/CFP ratio show the FRET response; g) and quantification (h) of FRET/CFP ratio of Mito-ABKAR in control and sg*SLC17A5* NRK cells reconstituted with Sialin (KRI/AAA), Sialin, and Sialin2. N = 30 cells from representative experiments of three repeats. i, Quantification of FRET/CFP ratio of Lyso-ExRai-ABKAR in control and sg*SLC17A5* NRK cells treated with metformin (Met, 10 mM) for 4 h. N = 30 cells from representative experiments of three repeats. See images in Extended Data Fig. 9h. j, k, PLA of Sialin/Sialin2-LKB1 in NRK (j) or HEK293T (k) cells treated with nitrate (4 mM, 4 h). N = 30 cells from representative experiments of three repeats. See images of HEK293T cells in Extended Data Fig. 10c. l, PLA of Sialin/Sialin2-CaMKK2 in HeLa cells treated with nitrate (4 mM, 4 h). N = 30 cells from representative experiments of three repeats. See images in Extended Data Fig. 10d. m, Quantification of FRET/CFP ratio of Mito-ABKAR in HeLa cells treated with nitrate (4 mM, 4 h). N = 30 cells from representative experiments of three repeats. See images in Extended Data Fig. 10e. n, o, IF staining images of LKB1 and MitoTracker in control and sg*SLC17A5* NRK cells reconstituted with Sialin (KRI/AAA), Sialin, and Sialin2 (n). Colocalization quantified by Pearson’s correlation coefficient (o). N = 30 cells from representative experiments of three repeats. p, Immunoblot analysis of pAMPK^T172^, AMPK, Flag, and Sialin2 in control and sg*SLC17A5* HEK293T cells reconstituted with vector, Flag-LKB1, Flag-LKB1-M, or Flag-LKB1-M (D194A). Representative images of n = 3 independent experiments were shown. q, r, IF staining images (upper, representative YFP images, lower, representative pseudocolor images of FRET/CFP ratio show the FRET response; q) and quantification (r) of FRET/CFP ratio of Mito-ABKAR in control and sg*SLC17A5* HEK293T cells reconstituted with vector, Flag-LKB1, Flag-LKB1-M, or Flag-LKB1-M (D194A). N = 30 cells from representative experiments of three repeats. s, Schematic illustration of nitrate enhances Sialin2-LKB1 interaction, promotes the recruitment of LKB1 to mitochondria, which in turn promotes the phosphorylation and activation of AMPK. For all panels, data are represented as mean ± SD, *P* value denotes *t*-test. Scale bar: 10 μm.

To delineate the spatial regulation of AMPK regulation, we analyzed mitochondrial and lysosomal compartments. *SLC17A5* knockout abolished mitochondrial pAMPK^T172^ colocalization and significantly reduced AMPK activation in mitochondrial fractions (Extended Data Fig. S9a–c). Real-time tracking using compartment-targeted FRET biosensors (Fig. 3e) revealed that nitrate specifically activated mitochondrial AMPK (Mito-ABKAR FRET/CFP ratio increase)^35^, an effect abolished in *SLC17A5* knockout cells (Fig. 3f, Extended Data Fig. S9d). Re-expression of wild-type Sialin2, but not the KRI/AAA mutant, fully restored nitrate-induced mitochondrial AMPK activation (Fig. 3g, h, Extended Data Fig. S9e, f). In contrast, treatment with metformin^36^, a lysosome- specific AMPK activator, induced comparable lysosomal AMPK activity in both control and *SLC17A5* knockout cells (Extended Data Fig. S9g). Similarly, glucose starvation significantly enhanced lysosomal AMPK activity, as measured using the lysosome-targeted biosensor Lyso-ExRai-AMPKAR, in both control and *SLC17A5* knockout cells (Fig. 3i, Extended Data Fig. S9h). These findings demonstrate that nitrate-Sialin2 signaling selectively regulates mitochondrial AMPK activation, without influencing lysosomal AMPK pools^37^.

Mechanistically, PLA and Co-IP revealed that nitrate enhanced the interaction between Sialin2 and LKB1 but not CaMKK2 (Fig. 3j–l, Extended Data Fig. S10a–d).

To dissect the kinase dependency of nitrate-induced AMPK activation, we utilized HeLa cells, which express CaMKK2 but lack endogenous LKB1^38^. In these cells, nitrate failed to activate AMPK, confirming an LKB1-dependent mechanism (Fig. 3m, Extended Data Fig. S10e). Further validation using the calcium ionophore A23187^36^, which activates CaMKK2, demonstrated comparable AMPK activation in both *SLC17A5* knockout and control HeLa cells (Extended Data Fig. S10f), reinforcing the specificity of Sialin2-LKB1 signaling. Critically, *SLC17A5* knockout disrupted nitrate- dependent recruitment of LKB1 to mitochondria, as evidenced by reduced colocalization and decreased mitochondrial LKB1 enrichment (Extended Data Fig. S11a–c). Wild-type Sialin2 fully rescued this defect but not the KRI/AAA mutant (Fig. 3n, o, Extended Data Fig. S11d), suggesting a coordinated translocation mechanism.

To directly establish the role of mitochondrial LKB1 localization in AMPK activation, we engineered mitochondrially targeted LKB1 (LKB1-M) and its kinase- dead mutant (LKB1-M D194A) (Extended Data Fig. S11e, f)^39,40^. In *SLC17A5* knockout cells, LKB1-M restored AMPK activation more effectively than cytosolic LKB1 (Fig. 3p–r). The kinase-dead LKB1-M mutant (D194A) partially rescued AMPK activation but was significantly less effective than wild-type LKB1-M (Fig. 3p–r), demonstrating that both mitochondrial localization and kinase activity are essential for nitrate-induced AMPK signaling.

Collectively, these findings define a nitrate-Sialin2-LKB1 axis that actively recruits LKB1 to mitochondria, driving compartmentalized AMPK activation (Fig. 3s). Unlike canonical AMPK activation mechanisms that respond to energy stress or calcium signaling, this pathway operates through the selective recruitment of LKB1 to mitochondria, enabling localized kinase activity without interfering with lysosomal AMPK pools, a paradigm of subcellular precision in metabolic adaptation.

### Nitrate-Sialin2 Signaling Promotes Mitochondrial Biogenesis and Functional Homeostasis via LKB1-AMPK Activation

Building on the discovery that nitrate-Sialin2 signaling activates AMPK via mitochondrial LKB1 recruitment, we investigated its functional impact on mitochondrial biogenesis and cellular homeostasis^41^. Immunoblot analysis revealed that nitrate treatment significantly increased the expression of N-terminal truncated PGC-1α (NT-PGC1α) and transcription factor A, mitochondrial (TFAM)—key regulators of mitochondrial biogenesis^42^, as well as mitochondrially encoded proteins, including mitochondrial NADH-ubiquinone oxidoreductase chain 5 (MT-ND5), cytochrome b (CYTB), mitochondrial cytochrome c oxidase II (MT-CO2), and ATP synthase F0 subunit 8 (ATP8) in control HEK293T cells (Fig. 4a, Extended Data Fig. S12a–d). These effects were abolished in *SLC17A5* knockout cells, and reintroduction of wild-type Sialin2, but not the KRI/AAA mutant, rescued nitrate-induced marker expression (Fig. 4a, Extended Data Fig. S12a–d). RT-qPCR analysis further confirmed the Sialin2-dependent upregulation of mitochondrial genes, including *TFAM*, *CYTB*, *NADH-ubiquinone oxidoreductase chain 1* (*ND1*), *ND5*, *MTCO2*, *mitochondrial cytochrome c oxidase III* (*MTCO3*), *ATP synthase F0 subunit 6* (*ATP6*), and *ATP8* (Extended Data Fig. S12e–l). IF staining for Cox IV, a mitochondrial complex IV subunit, demonstrated a Sialin2-dependent increase in mitochondrial mass upon nitrate treatment (Fig. 4b).

**Figure 4.**
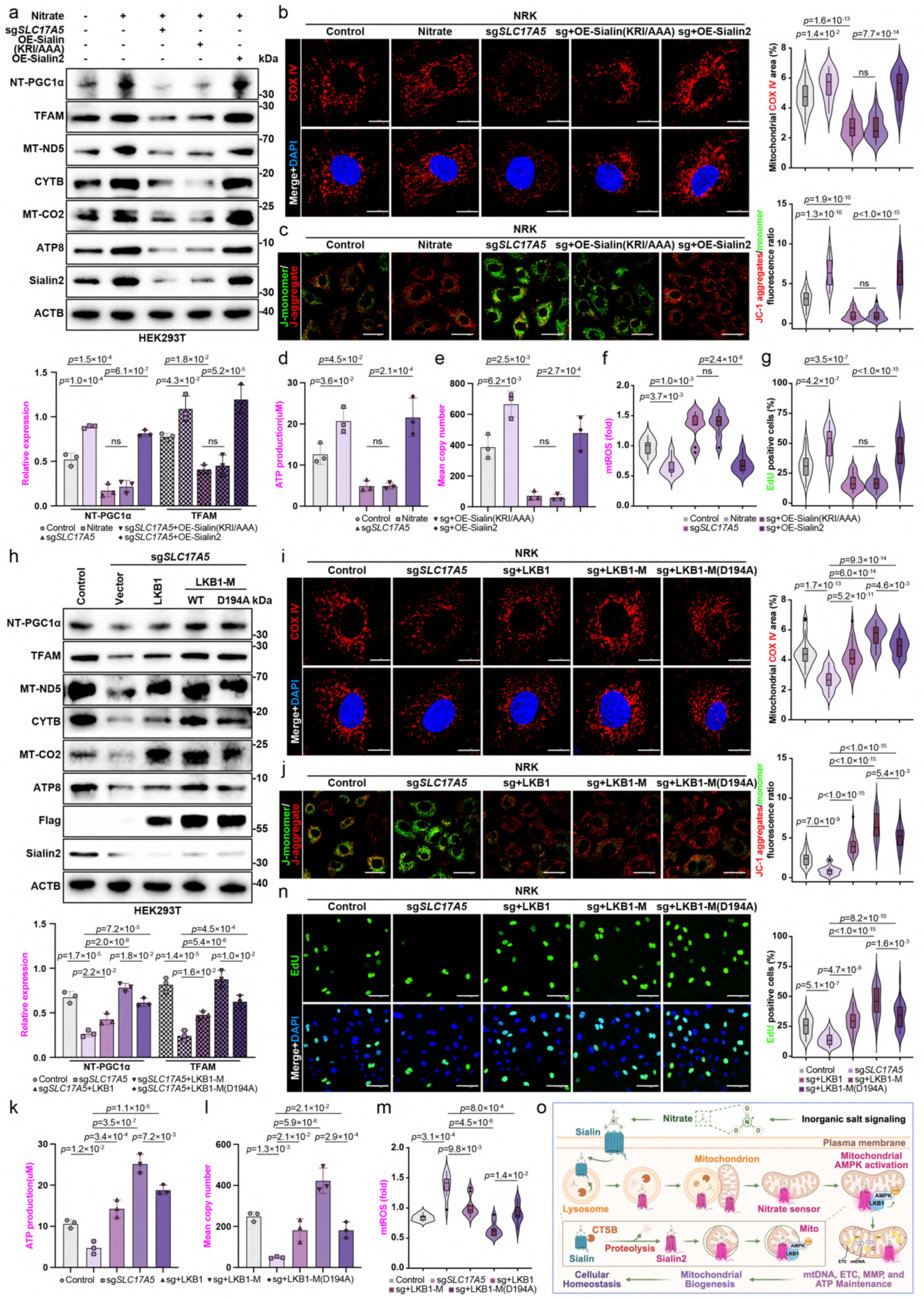
Nitrate-Sialin2 signaling promotes mitochondrial biogenesis and functional homeostasis via LKB1-AMPK activation a, Immunoblot analysis of NT-PGC1α, TFAM, mitochondrial-encoded proteins (MT- ND5, CYTB, MT-CO2, and ATP8), and Sialin2 in control and sg*SLC17A5* HEK293T cells reconstituted with Sialin (KRI/AAA) or Sialin2 and treated with nitrate (4 mM, 4 h). Representative images of n = 3 independent experiments were shown. See full quantitation in Extended Data Fig. 12a–d. b, c, IF staining images (left) and quantification (right) of Cox IV (mitochondria; b) and JC-1 (mitochondrial membrane potential; c) in control and sg*SLC17A5* NRK cells reconstituted with Sialin (KRI/AAA) or Sialin2 and treated with nitrate (4 mM, 4 h). N = 30 cells from representative experiments of three repeats. d, e, ATP production (d) and mitochondrial DNA (mtDNA) copy number (e) in control and sg*SLC17A5* NRK cells reconstituted with Sialin (KRI/AAA) or Sialin2 and treated with nitrate (4 mM, 4 h). N = 3. f, Mitochondrial ROS (mtROS) production in control and sg*SLC17A5* NRK cells reconstituted with Sialin (KRI/AAA) or Sialin2 and treated with nitrate (4 mM, 4 h). Production normalized to control. N = 3 from three independent experiments, each in triplicate. g, Proliferation (%EdU positive cells) in control and sg*SLC17A5* NRK cells reconstituted with Sialin (KRI/AAA) or Sialin2 and treated with nitrate (4 mM, 4 h). N = 30 cells from representative experiments of three repeats. See EdU images in Extended Data Fig. 12m. h, Immunoblot analysis of NT-PGC1α, TFAM, mitochondrial-encoded proteins (MT- ND5, CYTB, MT-CO2, and ATP8), Flag, and Sialin2 in control and sg*SLC17A5* HEK293T cells reconstituted with vector, Flag-LKB1, Flag-LKB1-M, or Flag-LKB1- M (D194A). Representative images of n = 3 independent experiments were shown. See full quantitation in Extended Data Fig. 13a–d. i, j, IF staining images (left) and quantification (right) of Cox IV (i) and JC-1 (j) in control and sg*SLC17A5* NRK cells reconstituted with vector, Flag-LKB1, Flag-LKB1- M, or Flag-LKB1-M (D194A). N = 30 cells from representative experiments of three repeats. k, l, ATP production (k) and mtDNA copy number (l) in control and sg*SLC17A5* NRK cells reconstituted with vector, Flag-LKB1, Flag-LKB1-M, or Flag-LKB1-M (D194A). N = 3. m, mtROS production in control and sg*SLC17A5* NRK cells reconstituted with vector, Flag-LKB1, Flag-LKB1-M, or Flag-LKB1-M (D194A). Production normalized to control. N = 3 from three independent experiments, each in triplicate. n, Proliferation (%EdU positive cells) in control and sg*SLC17A5* NRK cells reconstituted with vector, Flag-LKB1, Flag-LKB1-M, or Flag-LKB1-M (D194A). N = 30 cells from representative experiments of three repeats. o, Schematic illustration of nitrate triggers CTSB-mediated cleavage of Sialin at residues K256/R257/I258 to generate Sialin2, which translocates to mitochondria, acts as the nitrate sensor, recruits LKB1 to activate mitochondrial AMPK phosphorylation, and drives mitochondrial biogenesis to sustain mitochondrial function and cellular homeostasis. For all panels, data are represented as mean ± SD, *P* value denotes *t*-test. Scale bar: 10 μm.

To evaluate the functional consequences of Sialin2-mediated mitochondrial regulation, we quantified mitochondrial membrane potential, ATP production, mitochondrial DNA (mtDNA) copy number, and mitochondrial reactive oxygen species (mtROS) levels. Nitrate treatment significantly increased mitochondrial membrane potential, ATP synthesis, and mtDNA content while concurrently reducing mtROS accumulation in control cells (Fig. 4c–f, Extended Data Fig. S12q). These effects were abolished entirely in *SLC17A5* knockout cells but were rescued by wild-type Sialin2, whereas the KRI/AAA mutant failed to restore mitochondrial function (Fig. 4c–f, Extended Data Fig. S12q).

To directly assess whether LKB1-AMPK activation mediates Sialin2’s effects on mitochondrial biogenesis, we expressed mitochondrially targeted LKB1 (LKB1-M) and its kinase-dead mutant (LKB1-M D194A) in *SLC17A5* knockout cells. Strikingly, expression of LKB1-M, but neither cytosolic LKB1 WT nor the D194A mutant, restored nitrate-induced expression of NT-PGC1α, TFAM, and mitochondrial-encoded proteins/genes (Fig. 4h, Extended Data Fig. S13a–l). Likewise, LKB1-M rescued mitochondrial mass, membrane potential, ATP production, and mtDNA content while also suppressing mtROS accumulation (Fig. 4i–m, Extended Data Fig. S13p). These findings establish that both mitochondrial localization and kinase activity of LKB1 are indispensable for Sialin2-mediated mitochondrial biogenesis.

To assess the physiological relevance of this pathway, we performed cell viability and apoptosis assays under nitrate treatment. *SLC17A5* knockout cells exhibited reduced viability and increased apoptosis, phenotypes that wild-type Sialin2 rescued but not the KRI/AAA mutant (Fig. 4g, Extended Data Fig. S12m–o, r–t). Caspase-3 activity assays further confirmed that Sialin2 suppresses apoptosis, an effect abolished in *SLC17A5* knockout cells (Extended Data Fig. S12p, u). Notably, these pro-survival effects were exclusively rescued by LKB1-M, whereas cells expressing cytosolic LKB1 WT or the D194A mutant remained non-viable (Fig. 4n, Extended Data Fig. S13m–o, q–t).

Collectively, these results establish nitrate-Sialin2 signaling as a central regulator of mitochondrial biogenesis and metabolic homeostasis through LKB1-AMPK activation. Nitrate treatment enhances mitochondrial protein expression, mass, membrane potential, ATP synthesis, and cell survival, requiring Sialin2 and mitochondrial LKB1 kinase activity. These findings define a molecular axis linking extracellular nitrate levels to organelle-specific metabolic adaptation and cellular homeostasis (Fig. 4o)^43^.

## Discussion

Nitrogen homeostasis is essential for cellular physiology, yet the mechanisms by which mammalian cells directly sense nitrate (NO₃⁻), a critical inorganic nutrient and stress mitigator^18^, have remained elusive. Through a comprehensive integration of structural and functional analyses, we reveal that nitrate induces CTSB-mediated proteolytic cleavage of the nitrate transporter Sialin^14^, generating Sialin2. This mitochondria-targeted, high-affinity nitrate sensor orchestrates compartmentalized LKB1-AMPK signaling to regulate mitochondrial function (Extended Data Fig. S14). Importantly, our study reveals a unique negative feedback mechanism that dynamically regulates the dual roles of Sialin and Sialin2 in response to intracellular nitrate levels. When nitrate is scarce, cells upregulate full-length Sialin to enhance transmembrane nitrate import. As intracellular nitrate accumulates, Sialin undergoes specific proteolysis to generate Sialin2, which lacks the transmembrane domain and thus ceases further nitrate uptake. This structural shift not only terminates excess import but simultaneously enables Sialin2 to function as a nitrate sensor, interacting with signaling pathways to sustain homeostasis. When nitrate levels decline again, Sialin synthesis resumes—forming a tightly regulated “low-level transport activation—high-level sensing induction” closed-loop feedback that ensures dynamic equilibrium between metabolic demand and environmental input.

Notably, this proteolysis event produces two fragments: Sialin2 and an additional segment, which remains uncharacterized in terms of molecular identity and functional significance. For this study, we refer to this unidentified fragment as ’’Sialin-X’’, emphasizing its potential biological role awaiting further investigation. This is the first time we have identified a half transporter (Sialin2) that can exert specific cellular functions following protease digestion. This signaling axis directly connects extracellular nitrate levels to organelle-specific metabolic reprogramming, enhancing mitochondrial biogenesis, bioenergetic efficiency, and cell survival. Our findings redefine inorganic nitrate from a passive nitric oxide precursor to an autonomous signaling molecule that preserves mitochondrial integrity and systemic homeostasis, establishing a new paradigm of “inorganic salt signaling biology”.

Our study reveals a unique mammalian nitrate-sensing mechanism distinct from microbial and plant systems. Unlike bacterial Nar systems that regulate anaerobic respiration via transcription factors^44^, or plant nitrate transporter 1.1 (NRT1.1), which integrates dual-affinity transport with developmental signaling^45^, Sialin2 mediates a dual-function mechanism, simultaneous nitrate sensing and signal transduction, diverging from canonical nutrient sensors^46–48^. This strategy achieves remarkable specificity, evidenced by Sialin2’s high-affinity nitrate binding (*K*d = 1.08 ± 0.49 μM, Fig. 2i), which surpasses the low-affinity binding of plant NRT1.1^44^ and meets mammalian requirements for precision nitrate surveillance. By restricting AMPK activation to the mitochondrial compartment, Sialin2 enables localized metabolic adaptation without inducing deleterious global effects^49^, positioning it as a promising therapeutic target for mitochondrial disorders, including Alzheimer’s disease and metabolic syndrome.

However, several limitations warrant further exploration. While we can purify Sialin2 protein with good homogenity, the small size of the particle precludes the determination of high resolution cryo-EM structure, thus limiting the molecular mechanism of nitrate binding to Sialin2. Moreover, the functional nature of Sialin-X remains unknown—whether it retains partial activity, serves as a regulatory fragment, or undergoes further degradation requires additional investigation. Future studies should aim to characterize its biochemical properties and potential signaling roles. Although our biosensor (sCiSiNiS) confirms functional activation at physiologically relevant nitrate concentrations (10 μM, below the physiological plasma range of 20–50 μM^50^; Fig. 2m, Extended Data Fig. S6m), the *in vivo* threshold for sustained AMPK signaling requires further validation. In addition, potential crosstalk with stress- responsive pathways like mTOR-mediated anabolic regulation^51^ remains unexplored, leaving open questions about system-level integration of nitrate signaling.

Our findings suggest broader implications for inorganic salt signaling biology. The structural logic of Sialin2, with its ligand-specific conformational switch, may extend to other anions like sulfate or bicarbonate, expanding the repertoire of non-canonical signaling molecules. Furthermore, Sialin2 may facilitate inter-organelle communication beyond mitochondria or engage in retrograde nuclear signaling, potentially synergizing with classical nitric oxide pathways. Future studies should explore how inorganic signaling networks interface with canonical pathways to maintain systemic homeostasis, driving mechanistic discovery and therapeutic innovation.

## Methods

### Mice

C57BL/6 mice were used for all the experiments (The Jackson Laboratory). *Slc17a5*^−/−^ mice (on C57BL/6 mice background) were bred under specific-pathogen- free conditions in the animal facility of the Capital Medical University. Unless indicated otherwise, all other studies were performed with 8–10 weeks old male mice. The cages were kept at 18–24 °C ambient temperature under 40–60% humidity. The mice were maintained under a 12 hour (h)–12 h light-dark cycle from 06:00 to 18:00. Food and water were available ad libitum. Mice were euthanized by an injection of an overdose of sodium pentobarbital (150 mg/kg). Animal experiments were conducted according to the NIH’s Guide for the Care and Use of Laboratory Animals and approved by the Animal Care and Use Committee of Capital Medical University (AEEI-2023-067). The disease mice were successfully modeled, followed by a daily gavage of 200 μl in 8.4 μg/μl sodium nitrate for 1 month.

The 8-month-old B6.Cg-Tg (APPswe, PSEN1dE9) 85Dbo/Mmjax mouse (hereafter referred to as “APP/PS1”), a mouse model commonly used to study Alzheimer’s disease (AD)^52^, were randomly divided into the AD group, and the AD + Nitrate group, with 6 mice in each group. Six mice of the same age were used as the control group (WT).

Ionizing radiation (IR)-mice were irradiated under isoflurane anesthesia. The treatment position was supine, and mice were placed on the accelerator treatment bed before being exposed to a single 5-Gy dose of local ionizing radiation (6 MV photon energy at 300 cGy/minute; 21EX, Varian), which was directed at both submandibular gland regions, with a focus-skin distance of 100 cm^8^. IR-mice were randomly divided into the IR group, and the IR + Nitrate group, with 6 mice in each group. 6 mice that did not receive IR were the control group (WT).

The mice were fed with a high-fat diet (HFD) (Rodent Diet with 60% kcal% fat, Research Diets) for 8 weeks to induce obesity^6^, and were randomly divided into the OB group and the OB + Nitrate group, with 6 mice in each group. 6 mice with a normal diet were used as the control group (WT).

A model of postmenopausal osteoporosis was induced using excision of bilateral ovariectomy (OVX) in female mice^20^, which were randomly divided into the OP group and the OP + Nitrate group, with 6 mice in each group. 6 mice with only surgical operation without ligation and ovary removal were used as a control group (WT).

### Cell Culture and Constructs

HEK293T (#CRL-3216), NRK-52E (#CRL-1571), and HeLa (#CCL-2) cells were purchased from ATCC. All cells were cultured in DMEM (Corning, #10-013-CV) supplemented with 10% fetal bovine serum (FBS, Hyclone, #SH30910.03) and 1% penicillin/streptomycin (Gibco, #15140-122). For glucose starvation, cells were rinsed twice with PBS (Corning, #SH30256.01) and then incubated in glucose-free DMEM (Gibco, #11966-025) supplemented with 10% FBS. All cells were maintained at 37 °C with 5% CO2. The cell lines used in this study were routinely tested for mycoplasma contamination, and mycoplasma-negative cells were used. None of the cell lines used in this study is listed in the database of commonly misidentified cell lines maintained by ICLAC. Nitrate (Sigma-Aldrich, #221341) was dissolved in water and stored at 4 °C. Bafilomycin A1 (Baf-A1, MedChemExpress, #HY-100558), Chloroquine (CQ, MedChemExpress, #HY-17589A), MG-132 (MedChemExpress, #HY-13259), CA-074 (MedChemExpress, #HY-103350), Metformin (Met, MedChemExpress, #HY-B0627), AICAR (MedChemExpress, #HY-13417), and A-23187 (MedChemExpress, #HY- N6687) were dissolved in dimethyl sulfoxide (DMSO) and stored at −20 °C. Baf-A1, CQ, MG-132, CA-074, Met, AICAR, or A-23187 were directly added to cell culture media with an equal volume of vehicle (DMSO) added into control wells. For cell treatment that requires medium change, fresh media were pre-warmed overnight in an empty dish in the same incubator, which minimizes disturbance to cells caused by medium change.

### Antibodies

Monoclonal antibodies against CTSB (Santa Cruz Biotechnology, clone H-5, #sc- 365558), Tomm20 (Cell Signaling, clone D8T4N, #42406), LAMP1 (Invitrogen, clone LY1C6, #MA1-164), pAMPK^T172^ (Cell Signaling, clone 40H9, #2535), AMPK (Cell Signaling, clone D5A2, #5831), LKB1 (Santa Cruz Biotechnology, clone G-12, #sc- 374300), CAMKK2 (Cell Signaling, clone 6G9, #50049), Cox IV (Cell Signaling, clone 3E11, #4850), PGC1α (Santa Cruz Biotechnology, clone D-5, #sc-518025), HA- tag (Cell Signaling, clone C29F4, #3724; clone 6E2, #2367), Flag-tag (Cell Signaling, clone D6W5B, #14793; clone 9A3, #8146), GAPDH (Cell Signaling, clone 14C10, #2118), ACTB (Proteintech, clone 2D4H5, #66009), HSP90 (Cell Signaling, clone C45G5, #4877), and polyclonal antibodies against Sialin (Invitrogen, #PA5-30517), pAMPK^T172^ (Invitrogen, #PA5-37821), LKB1 (Proteintech, #10746), MT-ND5 (Proteintech, #55410), CYTB (Proteintech, #55090), MT-CO2 (Proteintech, #55070), ATP8 (Proteintech, #26723), TFAM (Proteintech, #22586), GFP-tag (Abcam, #ab290), and mCherry (Abcam, #167453) were used in this study.

For immunoblotting analyses, antibodies were diluted at a 1:1000 ratio except for ACTB (clone 2D4H5, 1:20000). For immunoprecipitation, antibody-conjugated agarose was purchased from Santa Cruz Biotechnology and Proteintech, including agarose-conjugated antibodies against anti-LKB1 (#sc-374334AC) and anti-FLAG (#sc-166355AC). For immunostaining analyses and proximity ligation assay (PLA), antibodies were diluted at a 1:100 ratio.

### Plasmids, Lentivirus, Transfection, and Stable Cell Line Construction

For stable expression of proteins in this study, all DNA sequences were of human origin unless otherwise specified. Site-directed mutagenesis (MCLAB) was employed to generate all mutants.

The cDNAs for HA-Sialin, HA-Sialin (K256A/R257A/I258A) were inserted into vector plasmid GV657 (pcDNA3.1-CMV-MCS-3flag-EF1A-zsGreen-sv40-puromycin) by BamHI and KpnI. The wild gene Sialin and Sialin (K256A/R257A/I258A) mutant have a HA tag at the N-terminus. The cDNAs for Sialin were fused into GV218 (pGC- FU-EGFP-IRES-Puromycin) through in-fusion reactions by BamHI and AgeI restriction enzymes.

The cDNA for Sialin2 was inserted into the lentivirus vector GV341 (pGC-FU- 3FLAG-SV40-puromycin-library) by restriction enzymes using AgeI/NheI and into the lentivirus vector GV287 (pGC-FU-3FLAG-SV40-EGFP) by restriction enzymes using AgeI. The Sialin2 (D69A/T72A/Q102A) cDNA through BamHI and NheI restriction enzymes insert lentivirus vector plasmid GV513 (pGC-FU-3FLAG-CBh-gcGFP- IRES-puromycin).

The cDNAs for Sialin-HA, Sialin (K256A/R257A/I258A)-HA, and mCherry- Sialin2 were all inserted into vector GV348 (pGC-FU-3FLAG-SV40-puromycin- library) by enzymatic digestion using AgeI and EcoRI. The fusion of mCherry with Sialin2 was by the following primers forward:5’- caaggaggtggaggatcaatggtgagcaagg-3’ and reverse: 5’- attgatcctccacctccttgtactccagcc-3’.

The cDNAs for N-mCitrine-Sialin2-mCitrine-C, N-mCitrine-Sialin2 (D69A)- mCitrine-C, N-mCitrine-Sialin2 (T72A)-mCitrine-C, N-mCitrine-Sialin2 (Q102A)- mCitrine-C, and N-mCitrine-Sialin2 (D69A/T72A/Q102A)-mCitrine-C were obtained from the cDNA library of Genechem (Shanghai, China). The split mCitrine was linked to Sialin2 gene in both C-terminal and N-terminal. The Plasmid vector GV712 (pcDNA3.1-puromycin) and gene sequence were digested by XbaI restriction enzymes and completed cloning through the in-fusion recombination method.

The cDNAs for LKB1-Flag, Mito-LKB1 (containing the COX8A mitochondrial targeting sequence), and its mutant Mito-LKB1 (D194A) through BamHI and HindIII restriction enzymes were inserted vector plasmid CV702 (pcDNA3.1-3FLAG- Puromycin).

Mito-ABKAR (organelle-targeting sequence: MAIQLRSLFP LALPGMLALLGWWWFFSRKKA, N-terminus; Addgene: #61509) and Lyso-ExRai- AMPKAR (organelle-targeting sequence: full length of lysosome-associated membrane protein 1 (LAMP1), N-terminus; Addgene: # 192449) were created by Jin Zhang lab. The two fusion genes, Mito-ABKAR and Lyso-ExRai-AMPKAR, were inserted into the plasmid vector GV394 (pcDNA3.1(+)) by BamHI and XhoI restriction enzymes.

The target U6-sgRNA (*SLC17A5*) *3 for *SLC17A5* (target seq: GGACCGCACGCCTCTTCTAC, CGAGACCTGGCCCGGAACGA, GAGTGTTGCGTTAGTGGATA) carrying the selected guide sequence was used for lentivirus packaging to generate knockout cell lines. The lentivirus vector plasmid CV279 (LV-sgCas9-P2A-puro) and U6-sgRNA (*SLC17A5*) *3 sequence were digested by KpnI and NheI restriction enzymes, and complete cloning through in-fusion recombination method. The infection rate of HEK293T cells by CRISPR viruses was remarkably high, and no cell death was observed after puromycin treatment of infected pools when all uninfected control cells died.

For recombinant protein production, the full-length human Sialin (UniProt: Q9NRA2) and Sialin2 (amino acid: 1–277) were cloned into the pCAG vector (Invitrogen) with an N-terminal FLAG tag. To create a Sialin-GFP and Sialin2-GFP fusion protein, we attached the GFP gene (Clontech) to the carboxy terminus of Sialin and Sialin2 via PCR.

All lentivirus and plasmids were purchased from Shanghai Genechem Co., Ltd. All vector construction methods were in-fusion recombination. The viral vector was transfected into 293T cells using Lipofectamine 2000 (Invitrogen) together with two helper plasmids, psPAX2, and pMD2.G. Infectious lentiviruses were harvested 72 h post-transfection, rapid centrifugation to remove cell debris, and then filtered through 0.45 μm cellulose acetate filters.

Except for transient expression of split N-mCitrine-Sialin2-mCitrine-C, split N- mCitrine-Sialin2 (D69A)-mCitrine-C, split N-mCitrine-Sialin2 (T72A)-mCitrine-C, split N-mCitrine-Sialin2 (Q102A)-mCitrine-C, split N-mCitrine-Sialin2 (D69A/T72A/Q102A)-mCitrine-C, Mito-ABKAR and Lyso-ExRai-AMPKAR, all other experiments were based on stable cell lines with genetic knockout and/or stable expression of proteins. For transient transfection, cells were transfected with plasmids using Lipofectamine 2000. The culture medium was refreshed after 6 h. Target cells were infected with lentiviruses for stable cell line generation and selected with puromycin (1–2 μg/ml) for 5–7 days. The primers used for plasmid constructions are listed in Supplementary Table 1. The synthesized DNA plasmid sequences are listed in Supplementary Table 2.

### Immunoprecipitation and Immunoblotting

Cells were removed from the medium after the indicated treatment, washed once with ice-cold PBS, and lysed in an ice-cold RIPA lysis buffer system (Santa Cruz Biotechnology, #sc-24948) with 1 mM Na3VO4, 5 mM NaF, and 1× protease inhibitor cocktail (Sigma-Aldrich, #P8340). The cell lysates were then sonicated at 15% amplitude for 15 s. After sonication, the cell lysates were incubated at 4 °C with continuous rotation for 1 h and subsequently centrifuged at 14,000 rpm for 10 minutes (min) to collect the supernatant. The BCA Pierce Protein assay (Thermo Fisher Scientific, #A65453) measured the protein concentration in the supernatant according to the manufacturer’s instructions. Equal amounts of protein were used for further analysis.

For immunoprecipitation, 0.5–1 mg of cell lysate protein was incubated with 20 µl of antibody-conjugated agarose (Santa Cruz Biotechnology) at 4 °C for 24 h. For endogenous Sialin/Sialin2 and LKB1, or FLAG-tagged Sialin2 immunoprecipitation, 0.5–1 mg of cell lysates were incubated with 20 µl anti-Sialin (Invitrogen, #PA5-30517), anti-LKB1 (Santa Cruz Biotechnology, #sc-374334AC), or anti-FLAG (Santa Cruz Biotechnology, #sc-166355AC) mouse monoclonal IgG antibody-conjugated agarose at 4 °C for 24 h. Normal immunoglobulin (IgG)-conjugated agarose was used as a negative control (Santa Cruz Biotechnology, #sc-2343). After washing five times with lysis buffer, the protein complex was eluted with SDS sample buffer.

For immunoblotting of immunoprecipitated complexes, horseradish peroxidase (HRP)-conjugated antibodies were used to avoid non-specific detection of immunoglobulin in the immunoprecipitated samples. HRP-conjugated LKB1 (Santa Cruz Biotechnology, #sc-374334HRP) and Flag (Proteintech, #HRP-66008) antibodies were purchased.

For immunoblotting, 5–20 µg of protein was loaded. Equal amounts of total proteins were separated on a 4–20% precast polyacrylamide gel (Bio-Rad, #4561096) and transferred onto 0.2 µm nitrocellulose membranes using the Trans-Blot Turbo system. The membrane was blocked with StartingBlock Blocking Buffer (Thermo Fisher Scientific, #37542) for 30 min at room temperature and then incubated with primary antibody at 4 °C overnight. The membrane was then washed and incubated with horseradish peroxidase-conjugated secondary antibodies at room temperature for 1 h. The membrane was washed and exposed with SuperSignal West Pico PLUS Chemiluminescent Substrate (Thermo Fisher Scientific, #34577) in a ChemiDoc Imaging System (Bio-Rad). The intensity of protein bands was quantified using ImageJ. The unsaturated exposure of immunoblot images was used for quantification with the appropriate loading controls as standards. Statistical data analysis was performed with Microsoft Excel, using data from at least three independent experiments.

### Immunofluorescence (IF) and Confocal Microscopy

For immunofluorescence studies, cells were grown on a glass bottom dish (Cellvis, #D35C4-20-1.5-N). Cells were fixed with 4% paraformaldehyde (PFA) (Santa Cruz Biotechnology, #sc-281692) for 20 min at room temperature, then washed three times with PBS. Next, the cells were permeabilized with 0.1% Triton X-100 for 20 min and rewashed three times with PBS. The cells were then blocked with 3% BSA in PBS for 1 h at room temperature. After blocking, cells were incubated with a primary antibody overnight at 4 °C. The cells were then washed three times with PBS and incubated with fluorescent-conjugated secondary antibodies (Invitrogen, #A21206, #A10042, #A21202, and #A10037) for 1 h at room temperature. After secondary antibody incubation, the cells were washed three times with PBS, and nuclei counterstained with 1 µg/ml 4’,6-diamidino-2-phenylindole (DAPI) (Invitrogen, #D3571) in PBS for 30 min at room temperature. The cells were subsequently washed three times with PBS and prepared for imaging. The images were taken using a Nikon AX confocal microscope and controlled by NIS-elements-Viewer software (Nikon). All images were acquired using the 100× objective lens (N.A. 1.4 oil). For quantification, the mean fluorescent intensity of channels in each cell was measured by ImageJ. The colocalization of double staining channels was quantified by ImageJ using Pearson’s correlation coefficient (Pearson’s r), ranging between 1 and –1. A value of 1 represents perfect correlation, 0 means no correlation, and –1 indicates perfect negative correlation. The quantitative graph was generated by GraphPad Prism. The images were processed using ImageJ.

### Live Cell Imaging

In HeLa cells, successfully and stably expressing EGFP-Sialin and EGFP-Sialin2, PK Mito Deep Red (Genvivo, #PKMDR-2) and LysoBrite^TM^ Red (AAT Bioquest, #22645) staining were performed according to the manufacturer’s instructions. The procedure for Hessian imaging was performed as described in ref. 53. A commercial structured illumination microscope (HiS-SIM) was used to acquire and reconstruct the cell images. To further improve the resolution and contrast of the reconstructed images, a sparse deconvolution was used according to the method in ref. 54.

In NRK cells successfully and stably expressing EGFP-Sialin and mCherry-Sialin2, LysoTracker (Beyotime, #C1046 and #C1047), MitoTracker (Beyotimeo, #C1035 and #C1048), ER Tracker (Beyotime, #C1041 and #C1042), and GolgiTracker (Beyotime, #C1043 and #C1045) staining were performed according to the manufacturer’s instructions. All cell images are confocal images acquired with a Nikon AX confocal microscope.

### Electron Microscopy (EM) with Immunogold Labelling Analysis

Cells were fixed for 2 h in 0.1 M sodium phosphate buffer (pH 7.4) containing fresh 4% paraformaldehyde and 0.2% glutaraldehyde, rinsed in PBS, and then either processed with osmium fixation and plastic embedding (as described for regular EM). Plastic-embedded samples were cut into ∼80 nm sections using a Reichert Ultracut S microtome and transferred to formvar-carbon-coated copper grids. Staining was performed at 22 °C. Sections were etched in saturated sodium m-periodate (Sigma- Aldrich) for 3 min, washed three times in water, blocked in PBT buffer (0.1% Triton X-100 and 1% BSA in PBS) for 30 min, incubated with rabbit anti-GFP (Abcam, #ab290) antibody and rabbit anti-mCherry (Abcam, #ab167453) antibody in PBT buffer at 4 °C overnight, washed four times in PBS, incubated with protein A-conjugated gold particles for 20 min, washed twice in PBS and four times in water, and stained with lead citrate for contrast. Grids with immunogold-labelled sections were imaged the same way as for regular EM. After being dried in the air, samples were observed with a transmission electron microscope (Hitachi, HT7700) at a voltage of 100 kV.

### Protein Expression and Purification

The plasmids of Sialin and Sialin2 with a Flag tag used for cell transfection were prepared using the GoldHi EndoFree Plasmid Maxi Kit (CWBIO). The HEK293F cells (Invitrogen) were cultured at 37 °C with 5% CO2 in a Multitron-Pro shaker (Infors) at a speed of 130 rpm. The plasmid was transiently transfected into the cells once the cell density reached 2.0 × 10^6^ cells/ml. For transfection of one liter of cell culture, approximately 1.5 mg of the plasmid was mixed with 3 mg of polyethylenimines (PEIs) (Polysciences) in 50 ml of fresh medium. The mixture was incubated for 15 min before adding it to the cell culture. 60 h later, Cells were collected by centrifugation at 4,000 *g* for 10 minutes after 60 h of transfection and resuspended in a buffer containing 25 mM HEPES (pH 7.5), 150 mM NaCl, and a mixture of three protease inhibitors: aprotinin (1.3 μg/ml, AMRESCO), pepstatin (0.7 μg/ml, AMRESCO), and leupeptin (5 μg/ml, AMRESCO).

For protein purification, the cells were incubated with 1.5% (w/v) n-dodecyl β-d- maltoside (DDM, Anatrace) at 4 °C for 2 h. Afterward, the cells were centrifuged at 18,000 *g* for 1 h to remove cell debris. The supernatant was loaded onto anti-FLAG M2 affinity resin (Sigma). The resin was washed with a wash buffer containing 25 mM HEPES (pH 7.5), 150 mM NaCl, and 0.02% GDN (w/v), followed by protein elution with the wash buffer plus 0.2 mg/ml FLAG peptide. The protein complex was then subjected to size-exclusion chromatography (Superose 6 Increase 10/300 GL, GE Healthcare) using a buffer containing 25 mM HEPES (pH 7.5), 150 mM NaCl, and 0.02% GDN. The peak fractions were collected for storage.

### Cryo-EM Sample Preparation and Data Collection

The purified Sialin and Sialin2 were concentrated to approximately 8 mg/ml before being applied to glow-discharged holey carbon grids (Quantifoil Au R1.2/1.3). The samples of purified Sialin and Sialin2 were incubated with 50 mM sodium nitrate for 30 min on ice before grid freezing. Aliquots of the protein sample (3.5 μl) were placed on the grids. The grids were then blotted for 3 s or 3.5 s and flash-frozen in liquid ethane cooled by liquid nitrogen using the Vitrobot (Mark IV, Thermo Fisher Scientific). The cryo grids were transferred to a Titan Krios microscope operating at 300 kV, equipped with a Gatan K3 Summit detector and GIF Quantum energy filter. Movie stacks were automatically collected using EPU software (Thermo Fisher Scientific) with a slit width of 20 eV on the energy filter. The defocus range was set from –1.4 µm to –1.8 µm in super-resolution mode at a nominal magnification of 81,000 ×. Each stack was exposed for 2.99 s, with an exposure time of 0.09 s per frame, resulting in 32 frames per stack. The total dose rate for each stack was approximately 50 e-/Å^2^. Subsequently, the stacks were motion-corrected using MotionCor2 1.1.0^55^. After motion correction, the movie stacks were binned 2-fold, resulting in a pixel size of 0.855 Å/pixel. Dose weighting was then applied to the data^56^. The defocus values were estimated with Gctf^57^.

### Image Processing, Model Building, and Refinement

The Cryo-EM structures of Sialin and Sialin2 were solved in cryoSPARC 3.3.1. Particles were automatically picked using cryoSPARC 3.3.1^58–61^. After 2D classification, the micrographs with good particles were selected, and these particles were subjected to several cycles of 2D classification, Ab-Initio Reconstruction, and multiple cycles of heterogeneous refinement without symmetry using cryoSPARC 3.3.1^62^. Refer to Extended Data Fig. S4 for details of data collection and processing. Predicted models of sialin2 were first obtained using Alphafold2^63^, which was manually adjusted based on the cryo-EM map with Coot 0.8.2^64^. Each transmembrane residue was manually checked, and the chemical properties were considered during model building.

### Structural Prediction of Sialin2 with Nitrate-binding Domain

Regarding the Sialin2-nitrate binding prediction, AI-Bind is a deep learning pipeline that combines network-derived learning strategies with unsupervised pre- trained node features to optimize the exploration of binding properties between novel proteins and ligands^31^. The pipeline is compatible with various neural architectures, including VecNet, Siamese model, and VAENet. For ligands, AI-Bind takes isomeric SMILES as input, capturing the structures of ligand molecules. AI-Bind considers a search space consisting of all drug molecules available in DrugBank and naturally occurring compounds in the Natural Compounds in Food Database (NCFD). It can be expanded by utilizing larger chemical libraries like PubChem. For proteins, AI-Bind uses amino acid sequences obtained from protein databases such as Protein Data Bank (PDB), Universal Protein knowledgebase (UniProt), and GeneCards.

### Microscale Thermophoresis (MST) Assay

The MST assay measured the binding affinity of purified recombinant proteins (GFP-Sialin and GFP-Sialin2) or cell lysates expressing their GFP fusion proteins to nitrate in vitro, as described previously^65,66^. The target proteins were fluorescently labeled by Monolith Protein Labeling Kit RED-NHS 2nd Generation (Nano Temper, #MO-L011) following the manufacturer’s instruction. The lysates of the target proteins were incubated with MST buffer (50 mM Tris-HCl, pH 7.8, 150 mM NaCl, 10 mM MgCl2, and 0.05% Tween-20) diluted 1.5-fold to provide an optimal level of fluorescence. GFP protein was used as the negative control. The target-ligand mixtures were loaded into Monolith NT.115 Series capillaries (Nano Temper, #MO-K022), and the MST traces were measured by Monolith NT.115 pico. The binding affinity was auto-generated by MO. Control v1.6 software. The dissociation constant (*K*d) from triplicate reads of measurements was calculated using MO. Affinity Analysis software (version 2.3).

### The Design of Nitrate Biosensor sCiSiNiS and sCiSiNiS-based Nitrate Confocal Imaging

Biosensor design is often empirical and best supported by three-dimensional structural information^67^. Although the protein structure of Sialin2 was initially obtained (Fig. 2g) and the structure of the nitrate-binding domain is predictable (Fig. 2n), how nitrate triggers the conformational change of Sialin2 remained to be elucidated. It is possible that the Sialin2 would be sufficiently flexible with the predicted nitrate-binding domain (69, 70, and 102 aa) sitting near the middle to facilitate the reconstitution of full mCitrine protein from N-mCitrine and mCitrine-C after nitrate binding. To record fluorescence images of sCiSiNiS, the excitation was set at 488 nm, and emission was collected at 525–545 nm. Images were collected at 2 h after nitrate induction. The dynamic fluorescence intensity values (Ratio) were calculated as (*F* - *F*0)/*F*0, where *F*0 represents the starting point value.

### Fluorescence Resonance Energy Transfer (FRET)

Cells with different treatments were plated on a glass-bottom plate (Cellvis) and transfected with Mito-ABKAR. AMPK activity at the mitochondria apparatus was measured 24 h later. To examine AMPK activity, the cells were maintained in DMEM (Gibco) containing 25 mM glucose for basal conditions before image acquisition. The images were acquired using a Leica DM16000B total internal reflection fluorescence microscope (Leica), equipped with a mercury lamp as laser power. The images were acquired using a BP420/10 excitation filter, a 440/520 dichroic mirror and two emission filters (BP472/30 for cyan fluorescent protein (CFP) and BP542/27 for YFP). The pseudocolor images of the FRET/CFP ratio showing the FRET response were obtained using the FRET Wizard of ImageJ.

Cells with different treatments were grown on a glass-bottom plate (Cellvis) and transfected with Lyso-ExRai-AMPKAR. After 24 h, cells were incubated with or without a glucose-free medium for another 2 h. Images were then acquired by Olympus FV-3000 confocal microscope (100× oil objective, NA = 1.45). Dual GFP excitation- ratio imaging was performed (405 nm excitation, detection wavelength is 465–565 nm; 488 nm excitation, detection wavelength is 465–565 nm). The fluorescence emission with 488 nm excitation and 405 nm excitation was measured using ImageJ software, and the ratio of these two fluorescence intensities (excitation 488/excitation 405) was calculated for further analysis.

### Proximity Ligation Assay (PLA)

PLA was utilized to detect in situ protein-protein/PI interaction as previously described. After fixation and permeabilization, cells with different treatments were blocked before incubation with primary antibodies, as in routine IF staining. The cells were processed by the PLA kit (MilliporeSigma, #DUO92101) according to the manufacturer’s instructions. The slides were then mounted with Duolink® In Situ Mounting Medium with DAPI (MilliporeSigma, #DUO82040). The Nikon AX confocal microscope detected PLA signals as discrete punctate foci and provided the intracellular localization of the complex. ImageJ was used to quantify the PLA foci.

### Mitochondrial Isolation

2 × 10^7^ HEK293T cells with different treatments were trypsinized and collected by centrifugation. The cell pellet was resuspended in 1.1 ml of hypotonic buffer containing 10 mM NaCl, 1.5 mM MgCl2, and 10 mM Tris (pH 7.5) and left on ice for 15 min. Swollen cells were next homogenized by Dounce homogenizer with 40 strokes of the tight pestle. Broken cells were added with 0.8 ml of 2.5 × homogenization buffer containing 525 mM D-mannitol, 175 mM sucrose, 2.5 mM EDTA, and 12.5 mM Tris (pH 7.5) and centrifuged at 1,300 *g* for 5 min twice. The resulting supernatant was then centrifuged at 12,000 *g* for 15 min. The final supernatant and pellet were regarded as cytosolic and mitochondria-enriched fractions, respectively. Mitochondria Isolation Kit (Thermo Fisher Scientific, #89874) was also used to purify the mitochondria fraction according to the manufacturer’s instructions.

### JC-1 Staining and Imaging

The JC-1 staining (Sigma-Aldrich, #CS0390) was used to measure mitochondrial membrane potential according to the vendor’s instructions. In brief, cells with different treatments were incubated with JC-1 (1.0 µg/ml) at 37 °C for 30 min. After three washes with PBS, changes in the ratio of JC-1 polymer (590 nm) to JC-1 monomer (520 nm) were used to indicate mitochondrial activity.

### Cellular ATP Measurement

ATP was determined using an ATP assay kit (Sigma-Aldrich, #MAK190). Briefly, ATP concentration is determined by phosphorylating glycerol, resulting in a colorimetric (570 nm) product proportional to the amount of ATP present. Lyse cells with different treatments in 100 ul of ATP assay buffer. Per the manufacturer’s instructions, measure the absorbance at 570 nm (A570) in a microplate reader.

### Quantification of the mtDNA Copy Number

Detection of mtDNA was carried out according to the protocol (TaKaRa, #7246). Total DNA was isolated from cells with the SpeeDNA Isolation Kit (ScienCell, #MB6918) according to the manufacturer’s instructions. DNA concentrations were measured using the Nanodrop 2000 (Thermo Fisher Scientific) and diluted to final concentrations of 20 ng/ml with double-distilled H2O. The mtDNA copy number was amplified using primers specific for the mitochondrial ND1, ND5, SLCO2B1, and SERPINA1 genes. Perform data analysis and determine the mtDNA copy number by relative quantification. Calculate the copy number of mtDNA from the Ct values obtained for each of the four target genes.

### Mitochondrial Superoxide (MitoSOX) Assay

Cells with different treatments were stained with MitoSO™ Red (Beyotime, #S0061) to detect intracellular mitochondrial superoxide. Cells were incubated in 5 μM working stock for 1 h, washed three times in warm PBS, and quantified by measuring the absorbance at 580 nm using a multifunctional fluorescent enzyme labeler.

### 5-Ethynyl-2’-deoxyuridine (EdU) Incorporation Assay

For the EdU incorporation assay, an EdU assay kit (Ribobio, #C10371) was used according to the manufacturer’s instructions. In brief, 2 × 10^4^ cells/well with different treatments were seeded into 24-well plates with coverslips and incubated overnight. Incubate for 2 h at 37 °C with medium containing EdU (final concentration, 50 μM). Subsequently, the cells were fixed with 4% formaldehyde for 30 min and treated with 0.5% Triton X-100 for 10 min for permeabilization. Cells were then administered an Apollo reaction cocktail for 30 min at room temperature and DNA stained with Hoechst 33342 for 30 min. The EdU incorporation rate was expressed as the ratio of EdU- positive cells to the total number of cells in each field.

### Crystal Violet Cell Viability Assay

In 96-well plates, 5 × 10^4^ cells/well with different treatments were seeded and incubated overnight. The cells were then fixed with 4% PFA and stained with 0.2% Crystal Violet. After washing the cells 3 times with PBS, the dye was extracted from the cells with 10% acetic acid and quantified by measuring the absorbance at 570 nm using a Synergy HTX Multi-Mode Microplate reader (BioTek Instruments, Inc.).

### Cell Death Detection ELISA

In 96-well plates, 5 × 10^4^ cells/well with different treatments were seeded and incubated overnight. Next, the cells were lysed and processed for cell death detection ELISA assay by following the manufacturer’s instructions (Sigma-Aldrich, #11774425001). At the end of the assay, the absorbance was read at 405 nm using a Synergy HTX Multi-Mode Microplate reader.

### Caspase-3 Activity Assay

In 6-well plates, 1 × 10^6^ cells/well with different treatments were seeded and incubated overnight. The cells were then lysed and processed for Caspase-3 activity by following the manufacturer’s instructions (Thermo Fisher Scientific, #E-13183). At the end of the assay, the fluorescence was read at excitation/emission 342/441 nm using a Synergy HTX Multi-Mode Microplate reader.

### RT-qPCR Assay

Total RNA was prepared from cells using TRIzol reagents (Invitrogen) according to the manufacturer’s instructions. RNA samples were reverse transcribed using the iScript cDNA Synthesis Kit (Bio-Rad) according to the provided protocol. The quantifications of gene transcripts were performed by qPCR using the CFX96 Touch Real-Time PCR detection system (Bio-Rad). GAPDH served as an internal control. A list of the PCR primers used to amplify the target genes is provided in Supplementary Table 3.

### Proteomic Assay

The sample was sonicated for three minutes on ice using a high-intensity ultrasonic processor (Scientz) in lysis buffer (1% SDS, 1% protease inhibitor cocktail, 1% phosphatase inhibitor). The remaining debris was removed by centrifugation at 12,000 *g* at 4 °C for 10 min. Finally, the supernatant was collected, and the protein concentration was determined with a BCA kit according to the manufacturer’s instructions. The protein sample was added with 1 volume of pre-cooled acetone, vortexed to mix, and added with 4 volumes of pre-cooled acetone, precipitated at -20 ℃ for 2 h. The protein sample was then redissolved in 200 mM TEAB and ultrasonically dispersed. Trypsin was added at 1:50 trypsin-to-protein mass ratio for the first digestion overnight. The sample was reduced with 5 mM dithiothreitol for 60 min at 37 °C and alkylated with 11 mM iodoacetamide for 45 min at room temperature in darkness. Finally, the peptides were desalted by the Strata-X SPE column.

Bio-material-based PTM enrichment (for phosphorylation): Peptide mixtures were first incubated with IMAC microspheres suspension with vibration in loading buffer (50% acetonitrile/0.5% acetic acid). To remove the non-specifically adsorbed peptides, the IMAC microspheres were washed sequentially with 50% acetonitrile/0.5% acetic acid and 30% acetonitrile/0.1% trifluoroacetic acid. To elute the enriched phosphopeptides, the elution buffer containing 10% NH4OH was added, and the enriched phosphopeptides were eluted by vibration. The supernatant containing phosphopeptides was collected and lyophilized for LC-MS/MS analysis.

The tryptic peptides were dissolved in solvent A and directly loaded onto a homemade reversed-phase analytical column (15 cm length, 100 μm i.d.). The separated peptides were analyzed in Orbitrap Astral with a nano-electrospray ion source. The electrospray voltage applied was 1900 V. Precursors were analyzed at the Orbitrap detector, and the fragments were analyzed at the Astral detector. The DIA data were processed using the Spectronaut (v18) search engine using software default parameters. The database was set as Homo_sapiens_9606_SP_20231220 (20429 entries); Trypsin/P was specified as a cleavage enzyme allowing up to 2 missing cleavages. The raw LC-MS datasets were first searched against the database and converted into matrices containing the Normalized intensity (the raw intensity after correcting the sample/batch effect) of proteins. GO annotation is to annotate and analyze the identified proteins with eggnog-mapper software (v2.0). The software is based on the EggNOG database. Extracting the GO ID from the results of each protein note, and then classifying the protein according to Cellular Component, Molecular Function, and Biological Process. We annotated the protein pathways based on the KEGG pathway database and performed BLAST alignments (blastp, evalue ≤ 1e-4) of the identified proteins. For the BLAST alignment results of each sequence, the alignment with the highest score (score) was selected to annotate.

### Statistics and Reproducibility

Statistical analyses were performed using GraphPad Prism 10 (GraphPad). An unpaired two-sided Student’s *t*-test was used to determine the significance between two groups of normally distributed data. Welch’s correction was used for groups with unequal variances. An ordinary one-way ANOVA was performed for multiple comparisons between groups, followed by Tukey’s or Dunnett’s test as specified in the legends. Brown-Forsythe and Welch’s correction was used for groups with unequal variances. *P* < 0.05 was considered as a statistically significant difference. The sample size was determined based on the previous studies and literature in the field using similar experimental paradigms. Each experiment was repeated at least three times independently, and the number of repeats is defined in each figure legend. We used at least three independent experiments or biologically independent samples for statistical analysis.

## Acknowledgments

S.W. is supported by grants from the Beijing Municipal Government grant (Beijing Laboratory of Oral Health, PXM2021_014226_000041 and PXM2021_014226_000020; Beijing Scholar Program-PXM2018_014226_000021), the National Natural Science Foundation of China (82030031 and 92149301), Chinese Research Unit of Tooth Development and Regeneration, Academy of Medical Sciences (No. 2019-12M-5-031). R.Y. is supported by grants from the Shenzhen Excellent Science and Technology Innovation Talent Training Program (RCYX20231211090342048) and the Major Talent Recruitment Program of Guangdong Province (2021QNO2Y167). X.L is supported by grants from the National Natural Science Foundation of China (82401082), the State Key Laboratory of Oral Diseases Open Fund (SKLOD2023OF11), Young Scientist Program of Beijing Stomatological Hospital, Capital Medical University (YSP202308). Fig. 1o, Fig. 2q, Fig. 3a, e, s, and Fig. 4o are generated using BioRender.

## Author contributions

S.W. and R.Y.: Conceived the project, acquired funding, provided direction, and supervised the research; X.L.: Coordinated the group, designed and conducted experiments, analyzed data, and interpreted results; S.Wu. and R.Y.: Screened the antibody and performed the structural experiments; O.J. and Z.C.: Assisted with cellular experiments; X.C., Y.F., and B.Z.: Helped the preparation of figure; Z.C., S.L., J.W., and J.Z.: Helped the preparation of experiments; G.X. and M.C.: Guided experimental design; X.L., R.Y., and S.W.: Wrote the manuscript, with input from all authors.

## Data availability

All data supporting the findings of this study are available in the article and its Supplementary Information section. Source data are provided in this paper.

## Extended Data Figures and Figure Legends Figure S1

## Supplementary Tables

**Figure S1.**
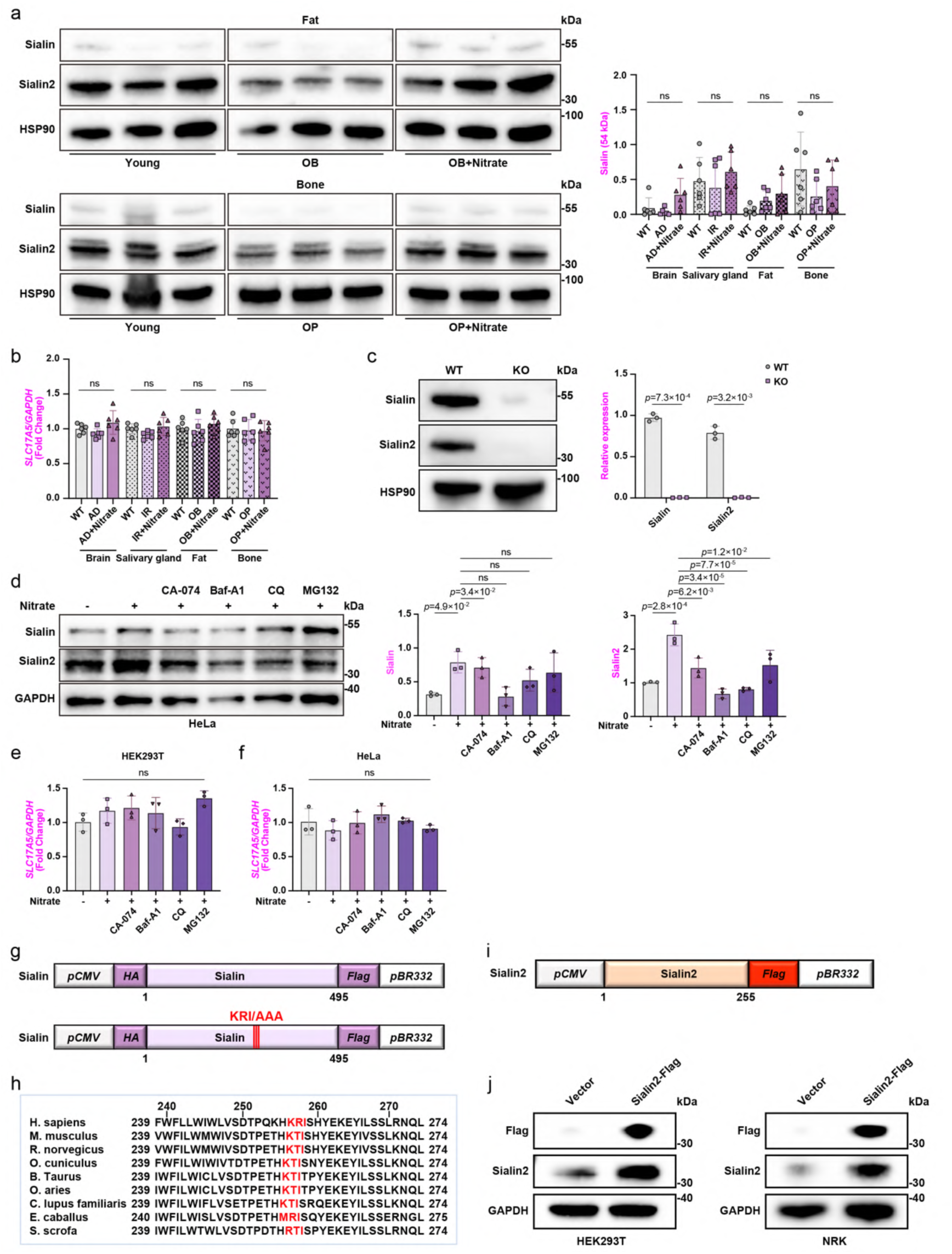
Characterization of Sialin and Sialin2 expression and processing in diverse models a, Immunoblot analysis of Sialin and Sialin2 protein normalized to HSP90 in white fat from obesity (OB) mice and bone from osteoporosis (OP) mice. Representative data of n = 6 independent experiments were shown. See full immunoblot images and quantitation in Fig. 1a. b, RT-qPCR analysis of *SLC17A5* mRNA. *SLC17A5* mRNA level was normalized to *GAPDH* mRNA. N = 6. c, Immunoblot analysis of Sialin and Sialin2 in cerebral cortex lysates from 3-week-old *Slc17a5* knockout (KO) mice. Representative images of n = 3 independent experiments were shown. d, Immunoblot analysis of Sialin and Sialin2 in HeLa cells pre-treated for 6 h with DMSO (vehicle), 20 μM CA-074 (CTSB inhibitor), 200 nM Baf-A1/40 μM CQ (lysosomal inhibitors), or 10 μM MG132 (proteasome inhibitor), followed by 4 h stimulation with 4 mM nitrate. Representative images of n = 3 independent experiments were shown. e, RT-qPCR analysis of *SLC17A5* mRNA in HEK293T cells. *SLC17A5* mRNA level was normalized to *GAPDH* mRNA. N = 3. f, RT-qPCR analysis of *SLC17A5* mRNA in HeLa cells. *SLC17A5* mRNA level was normalized to *GAPDH* mRNA. N = 3. g, Schematic of *HA-Sialin-Flag* gene and *HA-Sialin (KRI/AAA)-Flag* gene controlled by a human cytomegalovirus (CMV) promoter with the *pBR332* gene terminator. K, Lys; R, Arg; I, Ile. h, Key Sialin proteolytic sites (K256, R257, I258) were conserved in major mammals and outlined in red. i, Schematic of *Sialin2-Flag* gene controlled by a human cytomegalovirus (CMV) promoter with the *pBR332* gene terminator. j, Immunoblot analysis of Sialin2 and Flag in HEK293T (left panel) or NRK (right panel) cells stably expressed Sialin2-Flag. For all panels, data are represented as mean ± SD, *P* value denotes *t*-test.

**Figure S2.**
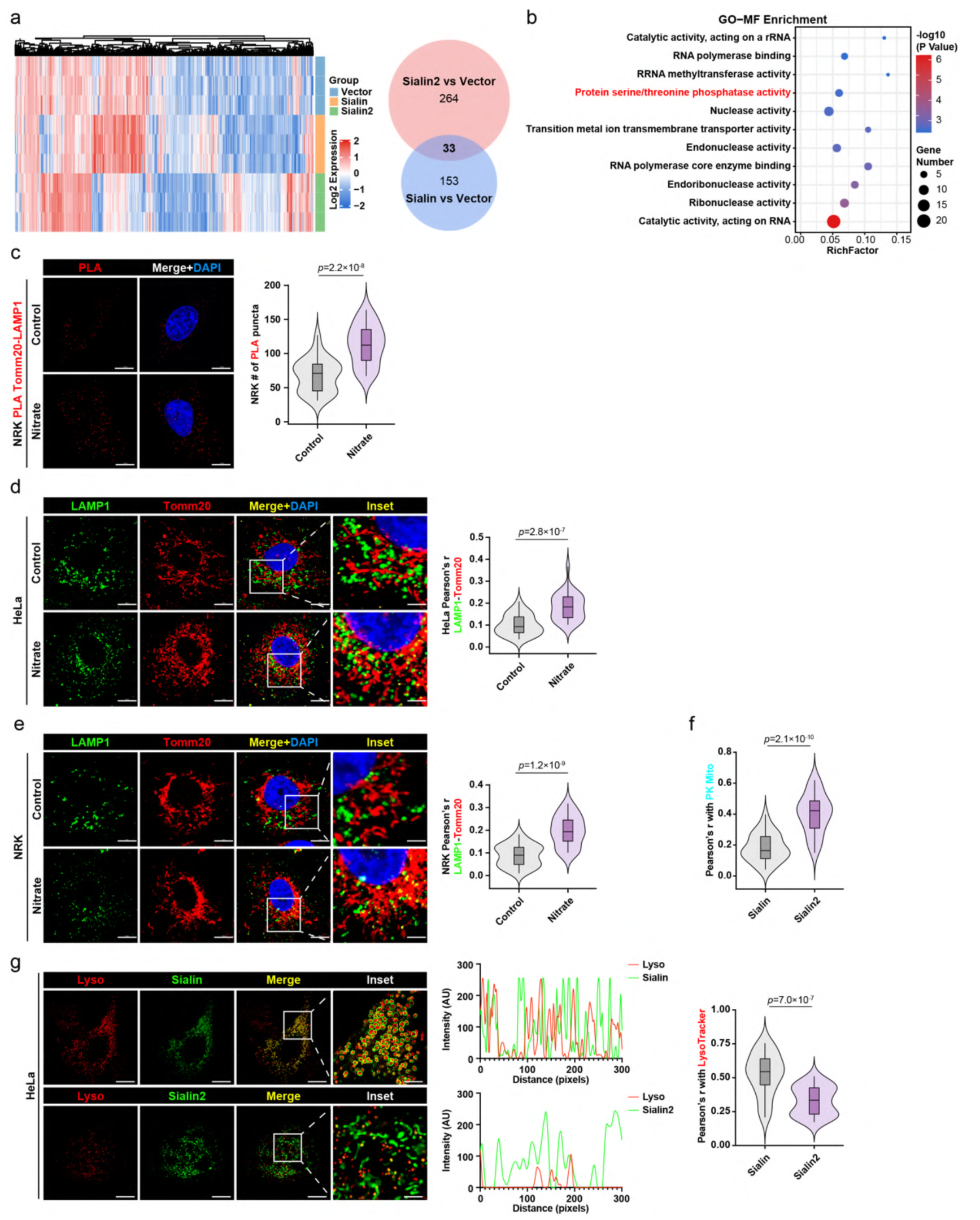
**Proteomic and subcellular localization analysis of Sialin and Sialin2** a, Heatmap and Venn diagram comparing proteomes associated with Sialin and Sialin2 in HEK293T cells. Data represented three biological replicates per condition. b, Top enriched cellular component GO terms for unique Sialin2-related proteomes in HEK293T cells listed by the rank of *P* value based on DAVID analysis. c, PLA of Tomm20 (mitochondria)-LAMP1 (lysosomes) in NRK cells treated with nitrate (4 mM, 4 h). N = 30 cells from representative experiments of three repeats. d, e, Immunofluorescence (IF) staining of Tomm20 and LAMP1 in HeLa (d) or NRK (e) cells treated with nitrate (4 mM, 4 h). Colocalization quantified by Pearson’s correlation coefficient. N = 30 cells from representative experiments of three repeats. f, Colocalization of Sialin or Sialin2 with mitochondria quantified by Pearson’s correlation coefficient. N = 30 cells from representative experiments of three repeats. See HiS-SIM live-cell images in Fig. 1m. g, HiS-SIM live-cell images of Sialin or Sialin2 colocalized with LysoTracker-labeled lysosomes in HeLa cells stably expressed GFP-Sialin or GFP-Sialin2. Colocalization quantified by Pearson’s correlation coefficient. N = 30 cells from representative experiments of three repeats. For all panels, data are represented as mean ± SD, *P* value denotes *t*-test.

**Figure S3.**
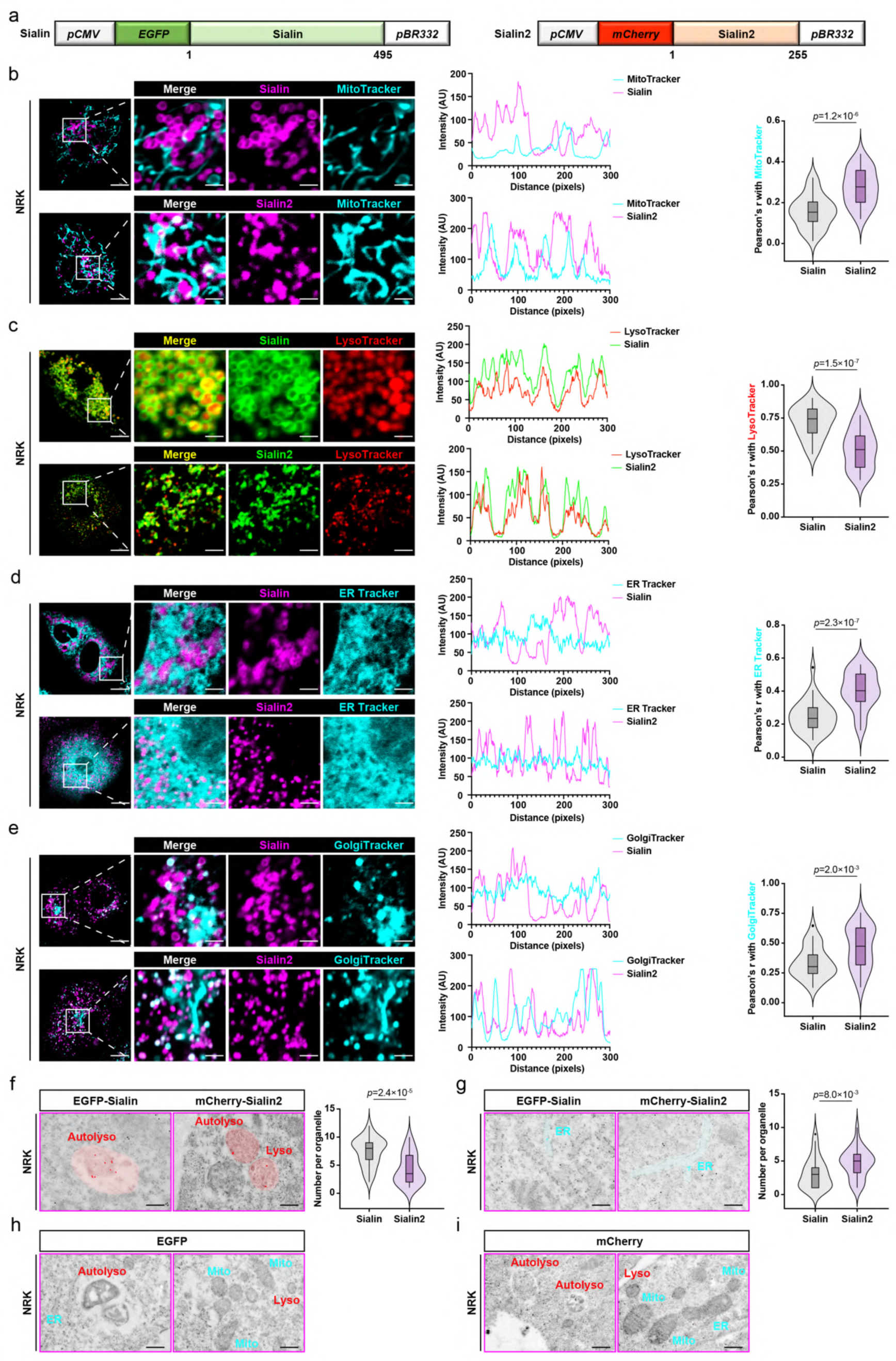
Subcellular localization of Sialin and Sialin2 in NRK cells a, Schematic of *GFP-Sialin* gene and *mCherry-Sialin2* gene controlled by a human cytomegalovirus (CMV) promoter with the *pBR332* gene terminator. b–e, Confocal live-cell images of Sialin or Sialin2 colocalized with MitoTracker- labeled mitochondria (b), LysoTracker-labeled lysosomes (c), ER Tracker-labeled endoplasmic reticulum (ER, d), and GolgiTracker-labeled Golgi apparatus (e) in NRK cells stably expressed GFP-Sialin or mCherry-Sialin2. Colocalization quantified by Pearson’s correlation coefficient. N = 30 cells from representative experiments of three repeats. f, g, Immunoelectron microscopy images of Sialin or Sialin2 localized to lysosomes (f) and ER (g) in NRK cells stably expressed GFP-Sialin or mCherry-Sialin2. N = 30 cells from representative experiments of three repeats. h, i, Immunoelectron microscopy images of GFP (h) or mCherry (i) in NRK cells stably expressed GFP-Sialin or mCherry-Sialin2. For all panels, data are represented as mean ± SD, *P* value denotes *t*-test.

**Figure S4.**
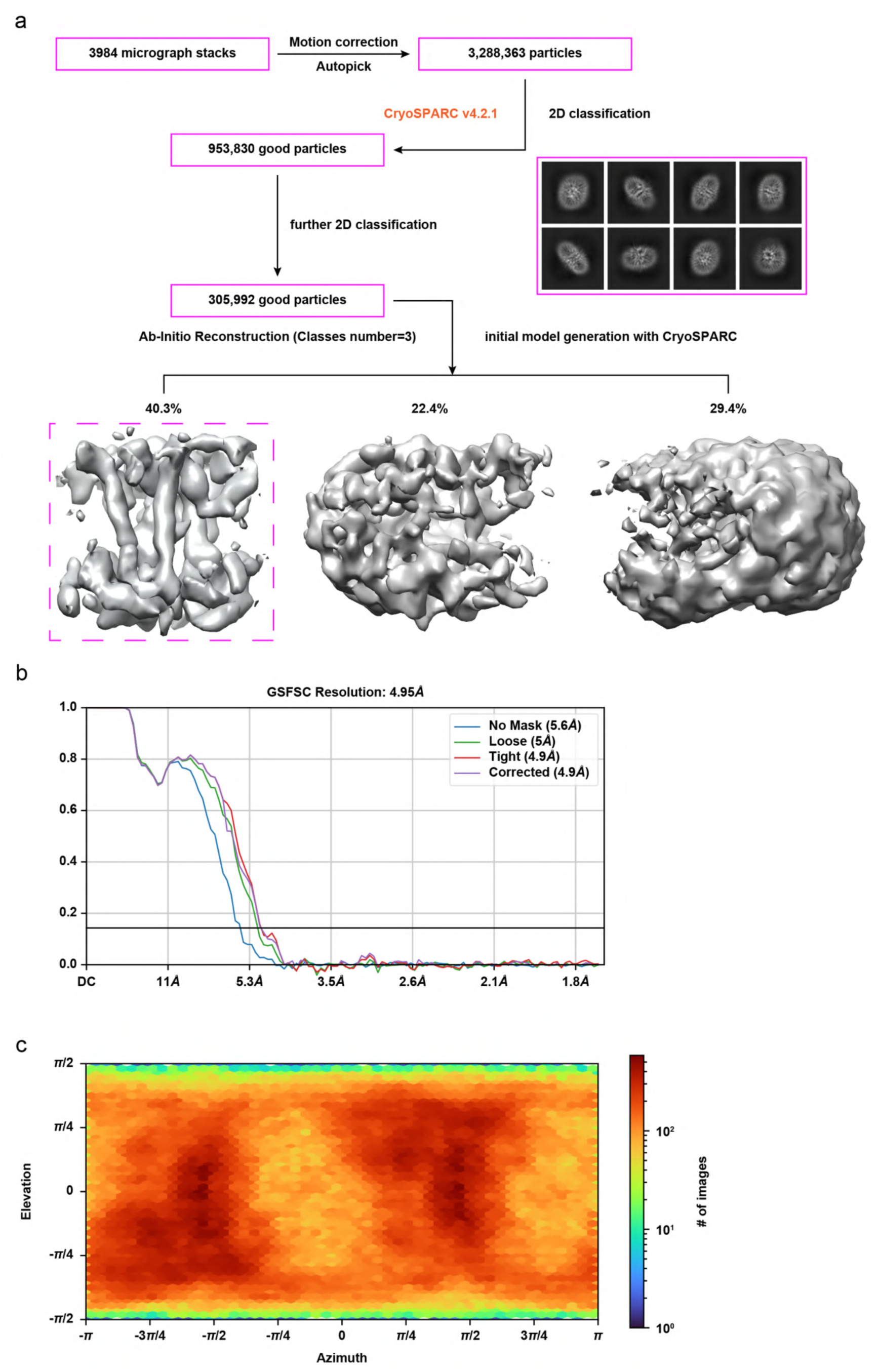
Flowchart of sialin2 for cryo-EM data processing a, Cryo-EM data processing workflow. The pipeline began with 3,984 micrographs, from which particles were selected through motion correction and template picking, yielding 3,288,363 initial particles. Following 2D classification, 953,830 high-quality particles were retained for ab-initio reconstruction. Subsequent refinement steps generated final 3D maps, revealing distinct particle populations at 22.4%, 29.4%, and 40.3% distributions. b, FSC (Fourier Shell Correlation) curve of the hetero-refined model of Sialin2. The curve represents the correlation between the refined model and the corresponding cryo- EM map, demonstrating the resolution and quality of the reconstruction. c, Angular distribution of the of 3D refinement using cryoSPARC 3.3.1.

**Figure S5.**
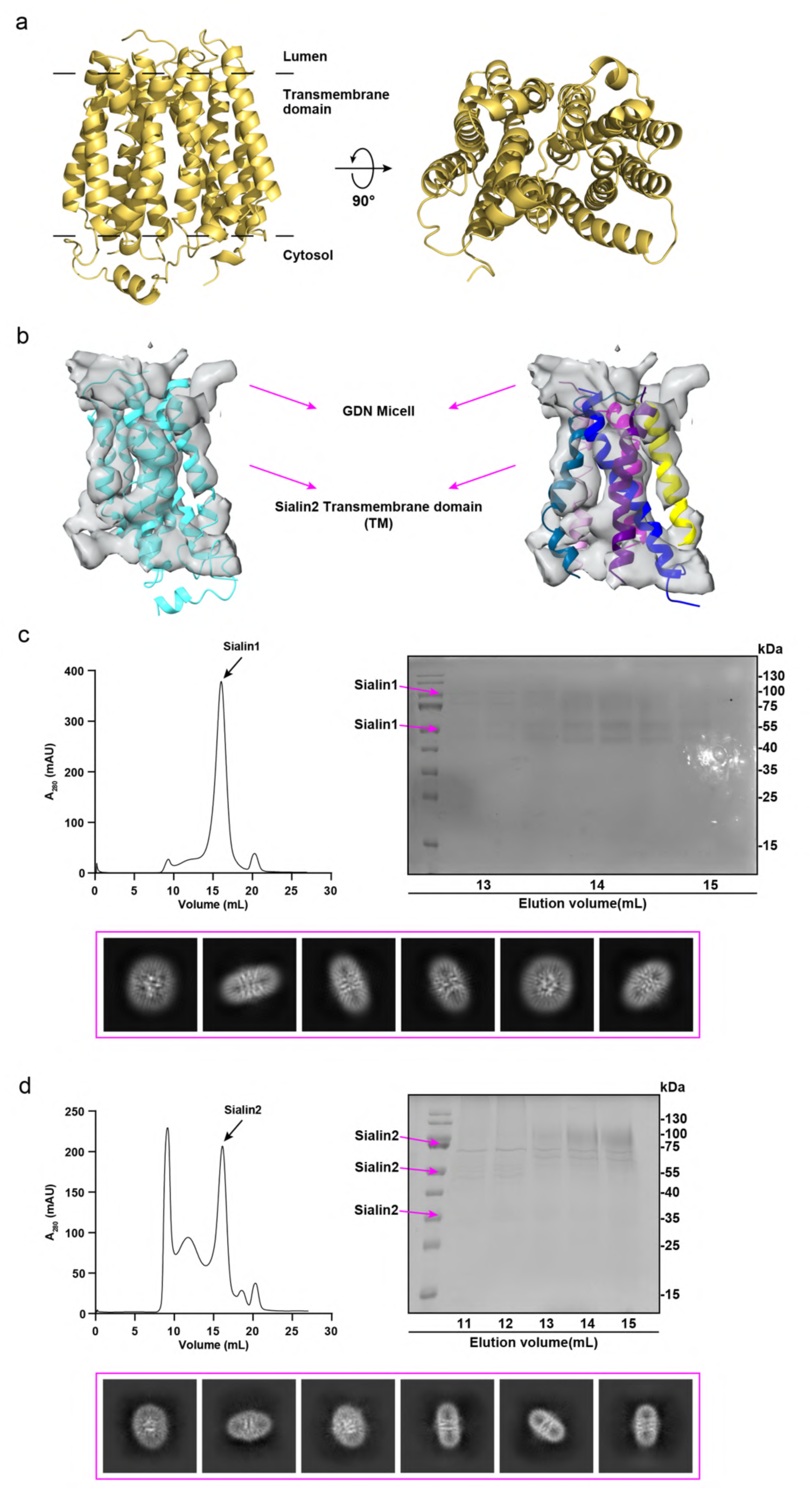
Biochemical characterization of the sialin and sialin2 a, Architecture of the Sialin Structure. The overall architecture of Sialin reveals a multi- domain organization. b, The left panel shows the overall cryo-EM map Sialin2 with predicted structure of Sialin2 while the right panel displays the map-fitting TM structure. c, d, Biochemical characterization and 2D-classificaton characterization of the Sialin and Sialin2, small multi-transmembrane domain proteins exhibit gradient multimerization patterns (35 kDa, 70 kDa, 105 kDa, etc.) on SDS-PAGE while maintaining homogeneous monomeric states in solution as 2D-classification results.

**Figure S6.**
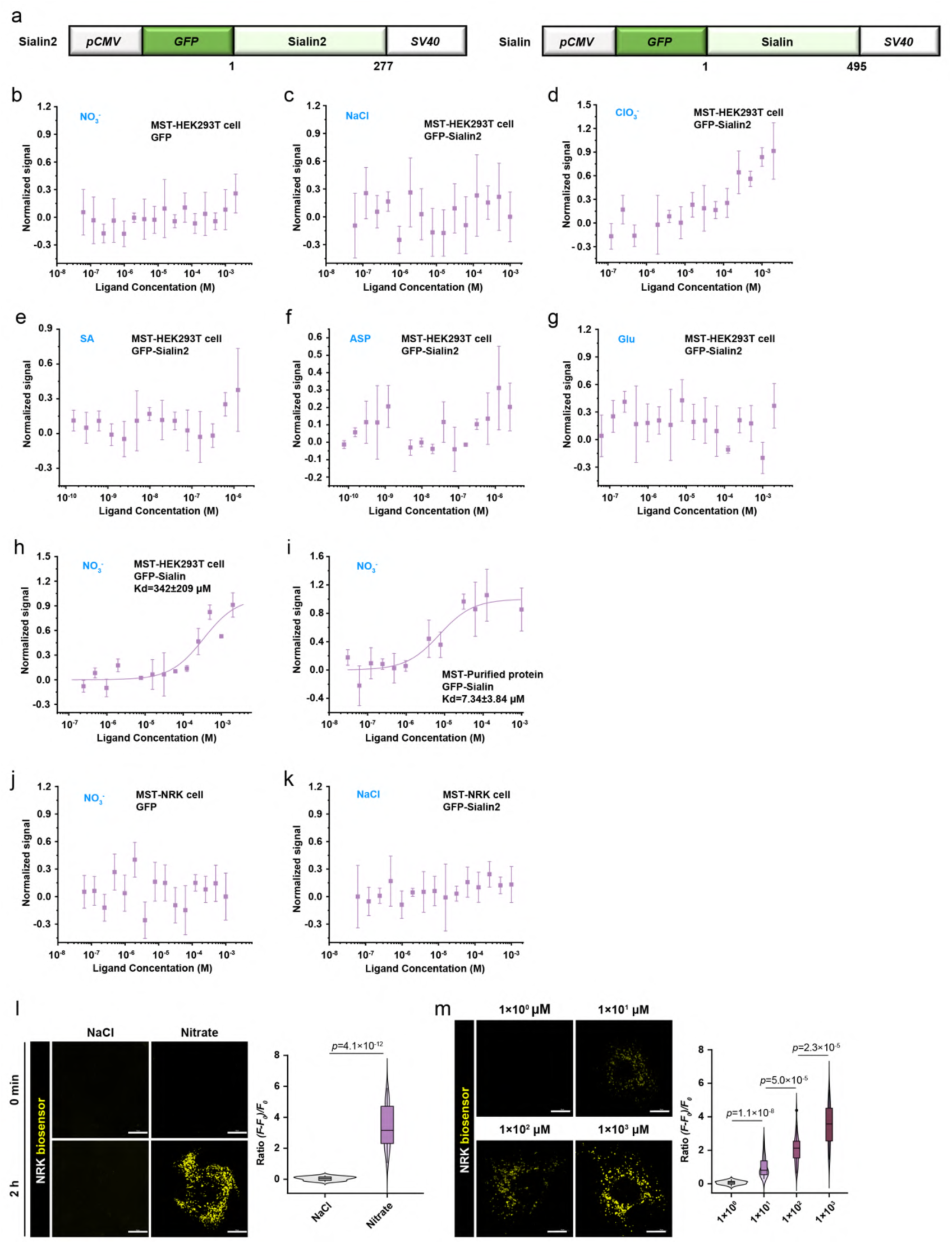
**Specificity and cross-cell validation of Sialin2 nitrate binding and biosensor function** a, Schematic of *GFP-Sialin* gene and *GFP-Sialin2* gene controlled by a human cytomegalovirus (CMV) promoter with the *SV40* gene terminator. b–e, MST assays confirmed no binding between nitrate and GFP control protein (b), NaCl and GFP-Sialin2 (c), ClO3^−^ and GFP-Sialin2 (d), sialic acid (SA) and GFP-Sialin2 (e), asparatic acid (ASP) and GFP-Sialin2 (f), or glutamic acid (Glu) and GFP-Sialin2 (g) in HEK293T cell lysate. h, i, Nitrate binding affinity (*K*d) of GFP-Sialin measured by MST in HEK293T cell lysate (342 ± 209 μM; h) and purified protein (7.34 ± 3.84 μM; i). N = 3. j, k, No binding detected between nitrate and GFP control protein (j) or NaCl and GFP- Sialin2 (k) in NRK lysate. l, m, Confocal live-cell imaging of cytoplasmic nitrate detected by sCiSiNiS in NRK cells treated with nitrate (4 mM; l) or dose-dependent nitrate (0–1000 μM; m) for 2 h. Fluorescence intensity normalized as (*F* - *F*0)/*F*0. N = 30 cells from representative experiments of three repeats. For all panels, data are represented as mean ± SD, *P* value denotes *t*-test. Scale bar: 10 μm.

**Figure S7.**
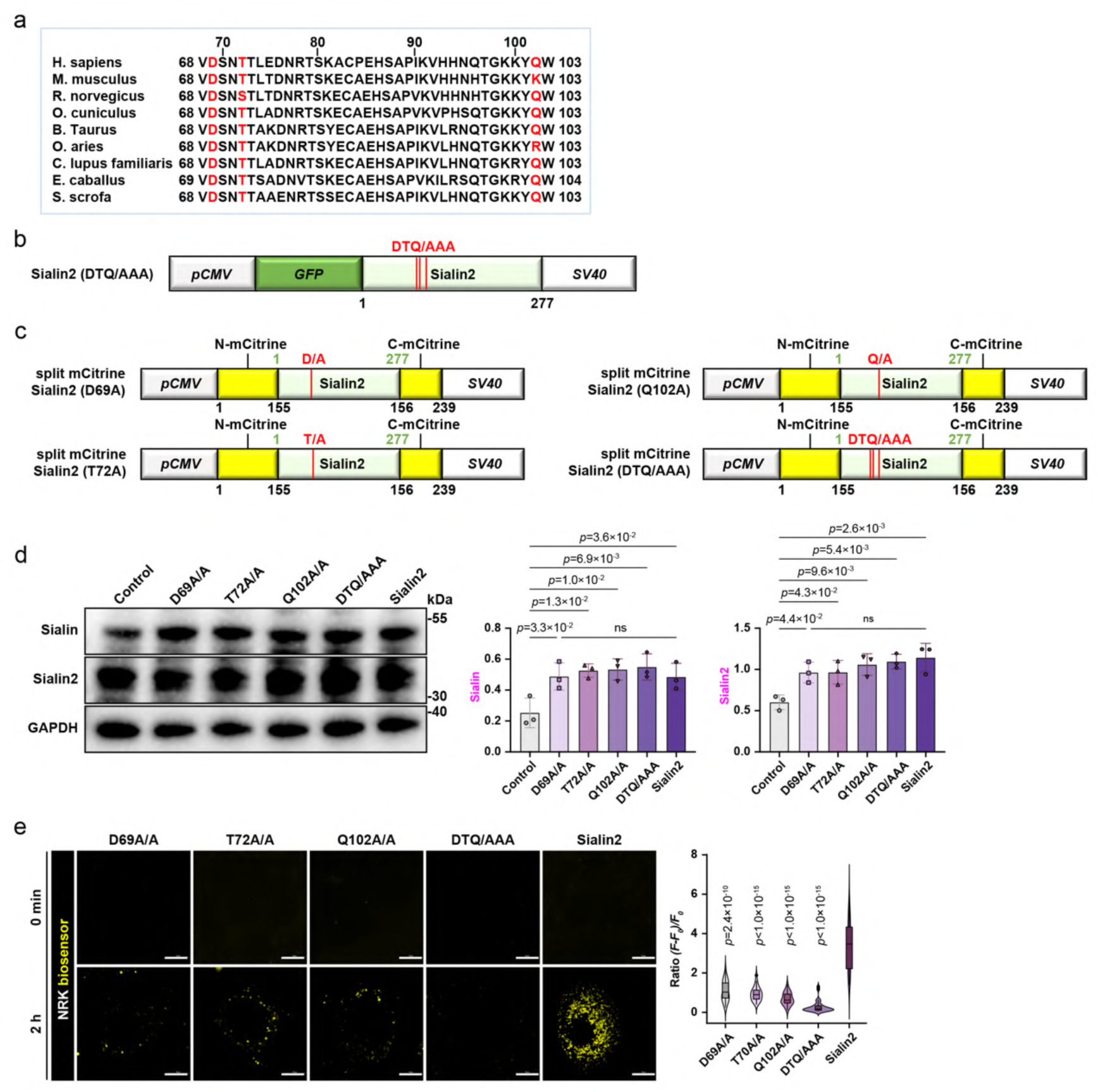
Mutational validation of nitrate-binding residues and reporter assays. a, Sequence alignment showed the conservation of critical nitrate-binding residues (red) in Sialin2 across major mammalian species. b, Schematic of mutant *GFP-Sialin2* (DTQ/AAA) gene controlled by a human cytomegalovirus (CMV) promoter with the *SV40* gene terminator. c, Structural models of sCiSiNiS variants with mutations in nitrate-binding residues (D69A, T72A, Q102A, or DTQ/AAA). d, Immunoblot analysis of Sialin and Sialin2 in HEK293T cells expressed sCiSiNiS, sCiSiNiS mutants (D69A, T72A, Q102A, or DTQ/AAA), or control vehicle. Representative images of n = 3 independent experiments were shown. e, Confocal live-cell imaging of sCiSiNiS mutants (D69A, T72A, Q102A, or DTQ/AAA) in NRK cells treated with nitrate (4 mM, 4 h). Fluorescence intensity normalized as (*F* - *F*0)/*F*0. N = 30 cells from representative experiments of three repeats. For all panels, data are represented as mean ± SD, *P* value denotes *t*-test. Scale bar: 10 μm.

**Figure S8.**
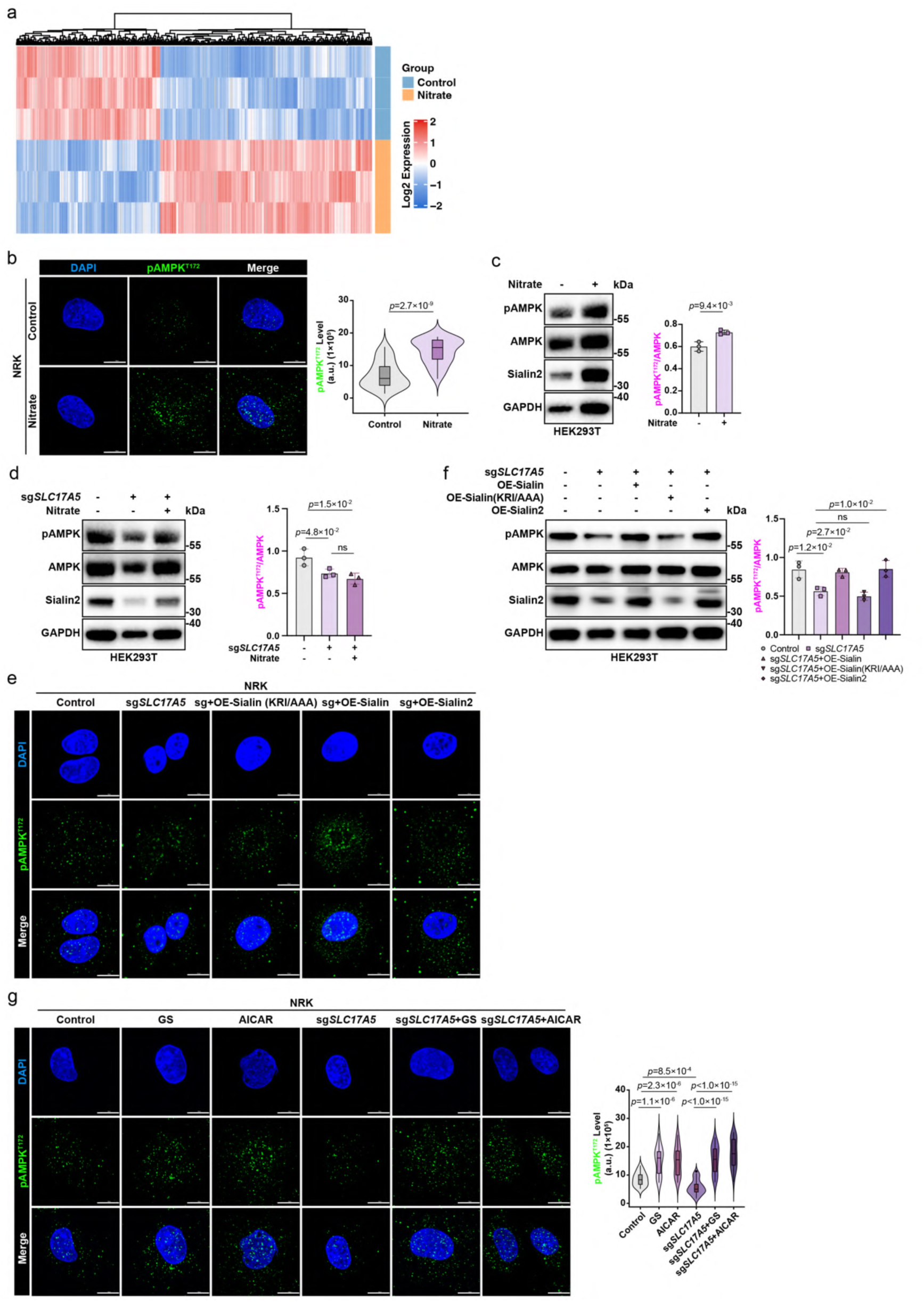
Nitrate-induced AMPK phosphorylation requires Sialin2 a, Heatmap showing differentially expressed (DE) genes (fold change ≥ 1.5, *P* ≤ 0.05) in HEK293T cells treated with nitrate (4 mM, 4 h). Data represent three biological replicates per condition. b, IF staining images (left) and quantification (right) of pAMPK^T172^ in NRK cells treated with nitrate (4 mM, 4 h). N = 30 cells from representative experiments of three repeats. c, Immunoblot analysis of pAMPK^T172^, AMPK, and Sialin2 in HEK293T cells treated with nitrate (4 mM, 4 h). Representative images of n = 3 independent experiments were shown. d, Immunoblot analysis of pAMPK^T172^, AMPK, and Sialin2 in control and sg*SLC17A5* HEK293T cells treated with nitrate (4 mM, 4 h). Representative images of n = 3 independent experiments were shown. e, IF staining images of pAMPK^T172^ in control and sg*SLC17A5* NRK cells reconstituted with Sialin (KRI/AAA), Sialin, and Sialin2. N = 30 cells from representative experiments of three repeats. See quantification in Fig. 3d. f, Immunoblot analysis of pAMPK^T172^, AMPK, and Sialin2 in control and sg*SLC17A5* HEK293T cells reconstituted with Sialin (KRI/AAA), Sialin, and Sialin2. Representative images of n = 3 independent experiments were shown. g, IF staining images (left) and quantification (right) of pAMPK^T172^ in control and sg*SLC17A5* NRK cells treated with glucose-free DMEM (GS, 2 h) or AICAR (2 mM, 2 h). Representative images of n = 3 independent experiments were shown. For all panels, data are represented as mean ± SD, *P* value denotes *t*-test. Scale bar: 10 μm.

**Figure S9.**
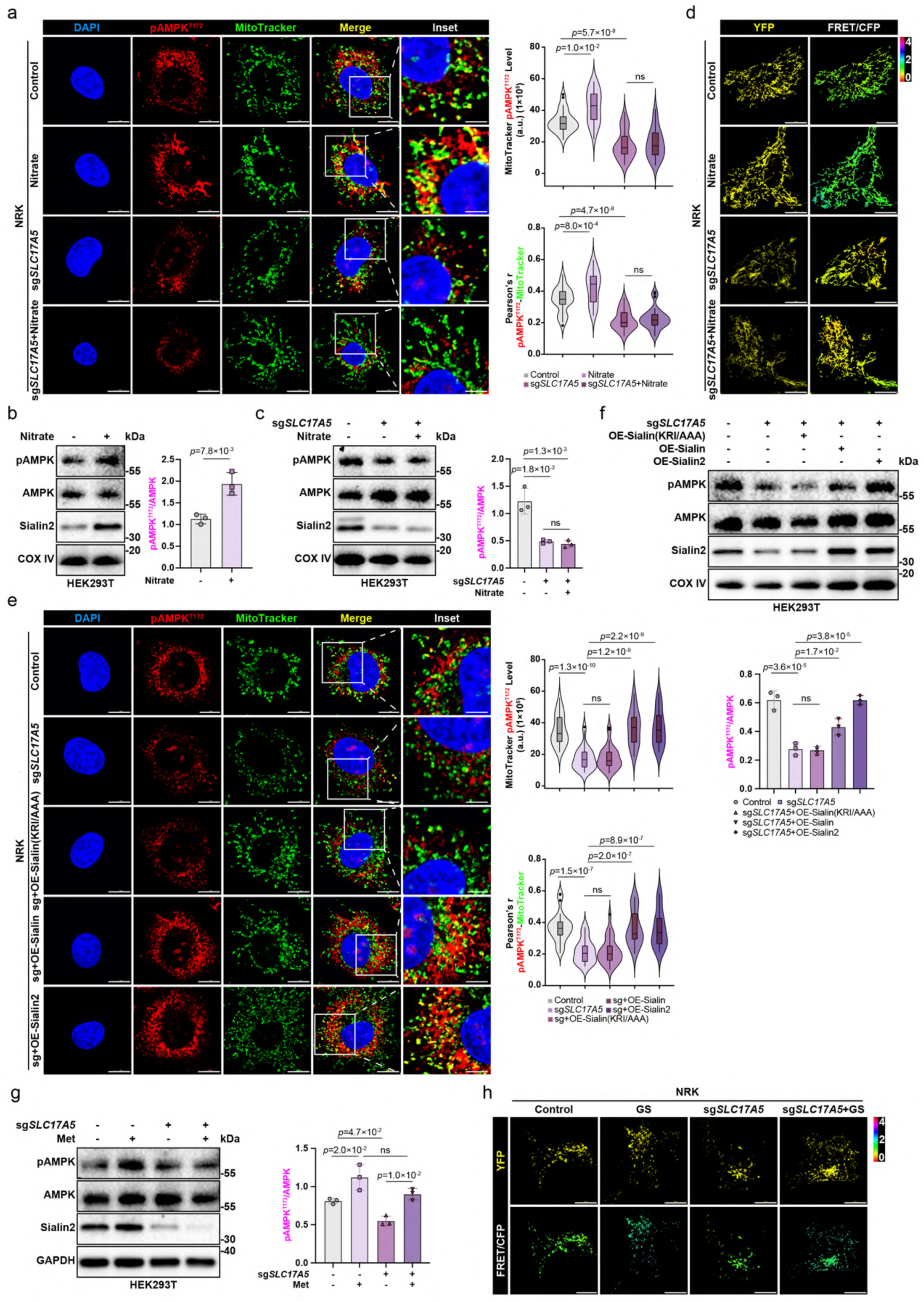
Nitrate-dependent AMPK phosphorylation and mitochondrial recruitment require Sialin2 a, IF staining images and quantification of pAMPK^T172^ and MitoTracker in control and sg*SLC17A5* NRK cells treated with nitrate (4 mM, 4 h). Colocalization quantified by Pearson’s correlation coefficient. N = 30 cells from representative experiments of three repeats. b, Immunoblot analysis of pAMPK^T172^, AMPK, and Sialin2 in mitochondrial fractions from HEK293T cells treated with nitrate (4 mM, 4 h). Representative images of n = 3 independent experiments were shown. c, Immunoblot analysis of pAMPK^T172^, AMPK, and Sialin2 in mitochondrial fractions from control and sg*SLC17A5* HEK293T cells treated with nitrate (4 mM, 4 h). Representative images of n = 3 independent experiments were shown. d, IF staining images (upper, representative YFP images, lower, representative pseudocolor images of FRET/CFP ratio show the FRET response) of FRET/CFP ratio of Mito-ABKAR in control and sg*SLC17A5* NRK cells treated with nitrate (4 mM, 4 h). N = 30 cells from representative experiments of three repeats. See quantification in Fig. 3f. e, IF staining images and quantification of pAMPK^T172^ and MitoTracker in control and sg*SLC17A5* HEK293T cells reconstituted with Sialin (KRI/AAA), Sialin, and Sialin2. Colocalization quantified by Pearson’s correlation coefficient. N = 30 cells from representative experiments of three repeats. f, Immunoblot analysis of pAMPK^T172^, AMPK, and Sialin2 in mitochondrial fractions from control and sg*SLC17A5* HEK293T cells reconstituted with Sialin (KRI/AAA), Sialin, and Sialin2. Representative images of n = 3 independent experiments were shown. g, Immunoblot analysis of pAMPK^T172^, AMPK, and Sialin2 in control and sg*SLC17A5* HEK293T cells treated with Met (10 mM, 4 h). Representative images of n = 3 independent experiments were shown. h, IF staining images (upper, representative YFP images, lower, representative pseudocolor images of FRET/CFP ratio show the FRET response) of FRET/CFP ratio of Lyso-ExRai-ABKAR in control and sg*SLC17A5* NRK cells treated with or without glucose-free DMEM (GS) for 2 h. N = 30 cells from representative experiments of three repeats. See quantification in Fig. 3i. For all panels, data are represented as mean ± SD, *P* value denotes *t*-test. Scale bar: 10 μm.

**Figure S10.**
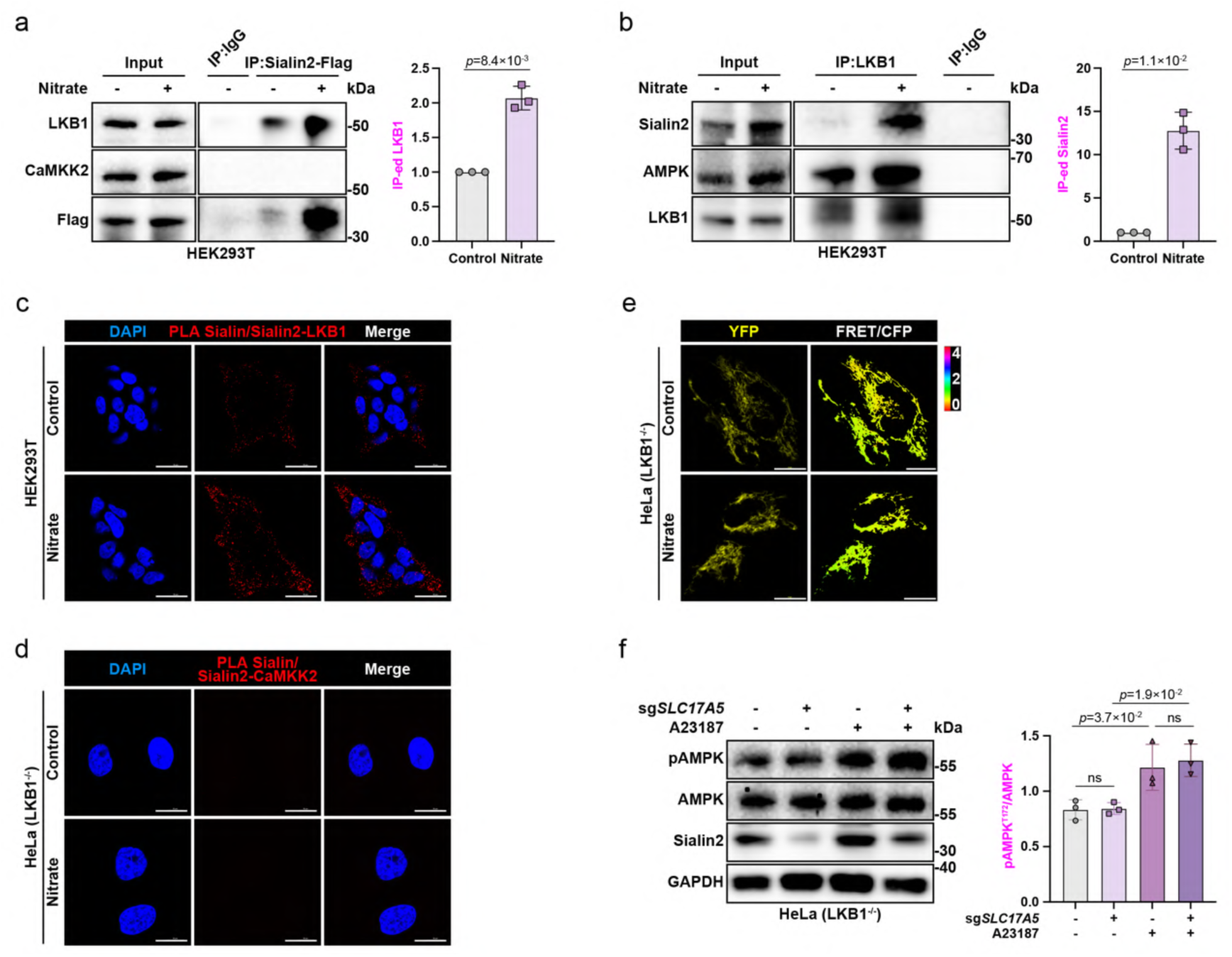
**Nitrate-stimulated Sialin2 specifically interacts with LKB1 to activate mitochondrial not lysosomal AMPK signaling** a, b, Co-IP of Sialin2 with LKB1/CaMKK2 (a) and LKB1 with Sialin2/AMPK (b) in HEK293T cells treated with nitrate (4 mM, 4 h). Representative data of n = 3 independent experiments were shown. c, d, PLA of Sialin/Sialin2-LKB1 in HEK293T cells (c) and Sialin/Sialin2-CaMKK2 in HeLa cells (d) treated with nitrate (4 mM, 4 h). N = 30 cells from representative experiments of three repeats. See quantification of HEK293T cells in Fig. 3k and quantification of HeLa cells in Fig. 3l. e, IF staining images (upper, representative YFP images, lower, representative pseudocolor images of FRET/CFP ratio show the FRET response) of FRET/CFP ratio of Mito-ABKAR in HeLa cells treated with nitrate (4 mM, 4 h). N = 30 cells from representative experiments of three repeats. See quantification in Fig. 3m. f, Immunoblot analysis of pAMPK^T172^, AMPK, and Sialin2 in control and sg*SLC17A5* HeLa cells treated with A23187 (2 μM, 30 min). Representative images of n = 3 independent experiments were shown. For all panels, data are represented as mean ± SD, *P* value denotes *t*-test. Scale bar: 10 μm.

**Figure S11.**
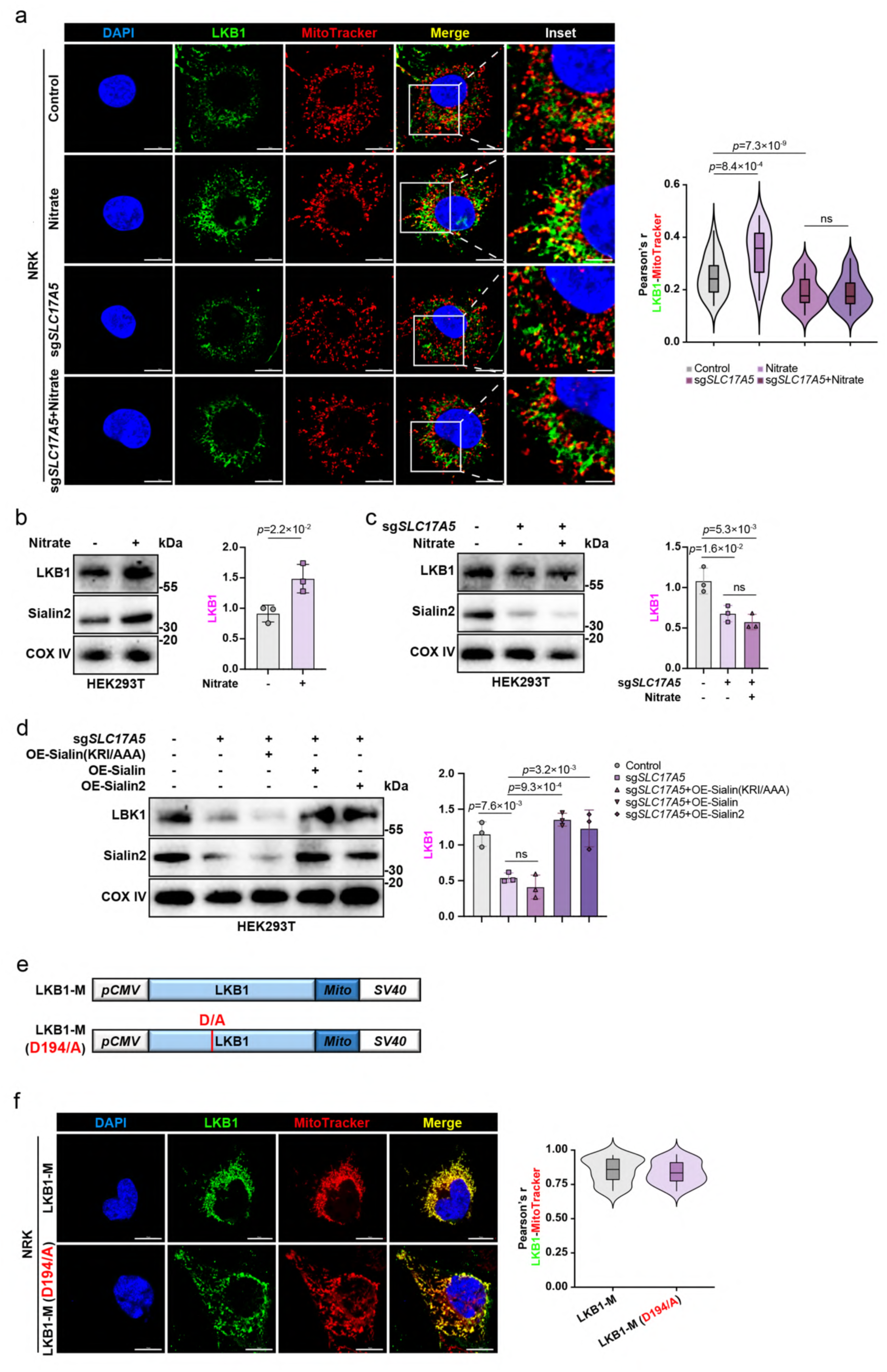
Sialin2-dependent mitochondrial recruitment of LKB1 is required for AMPK activation a, IF staining images of LKB1 and MitoTracker in control and sg*SLC17A5* NRK cells treated with nitrate (4 mM, 4 h). Colocalization quantified by Pearson’s correlation coefficient. N = 30 cells from representative experiments of three repeats. b, Immunoblot analysis of LKB1 and Sialin2 in mitochondrial fractions from HEK293T cells treated with nitrate (4 mM, 4 h). Representative images of n = 3 independent experiments were shown. c, Immunoblot analysis of LKB1 and Sialin2 in mitochondrial fractions from control and sg*SLC17A5* HEK293T cells treated with nitrate (4 mM, 4 h). Representative images of n = 3 independent experiments were shown. d, Immunoblot analysis of LKB1 and Sialin2 in mitochondrial fractions from control and sg*SLC17A5* HEK293T cells reconstituted with Sialin (KRI/AAA), Sialin, and Sialin2. Representative images of n = 3 independent experiments were shown. e, Schematics of the LKB1-COX8A (LKB1-M) and LKB1-M (D194A) chimaeras. The chimaeras contain full length of LKB1 or LKB1 D194A mutant and aa1-25 of COX8A. f, IF staining images showed the mitochondrial localization of LKB1-M and LKB1-M (D194A) in NRK cells. Colocalization quantified by Pearson’s correlation coefficient. N = 30 cells from representative experiments of three repeats. For all panels, data are represented as mean ± SD, *P* value denotes *t*-test. Scale bar: 10 μm.

**Figure S12.**
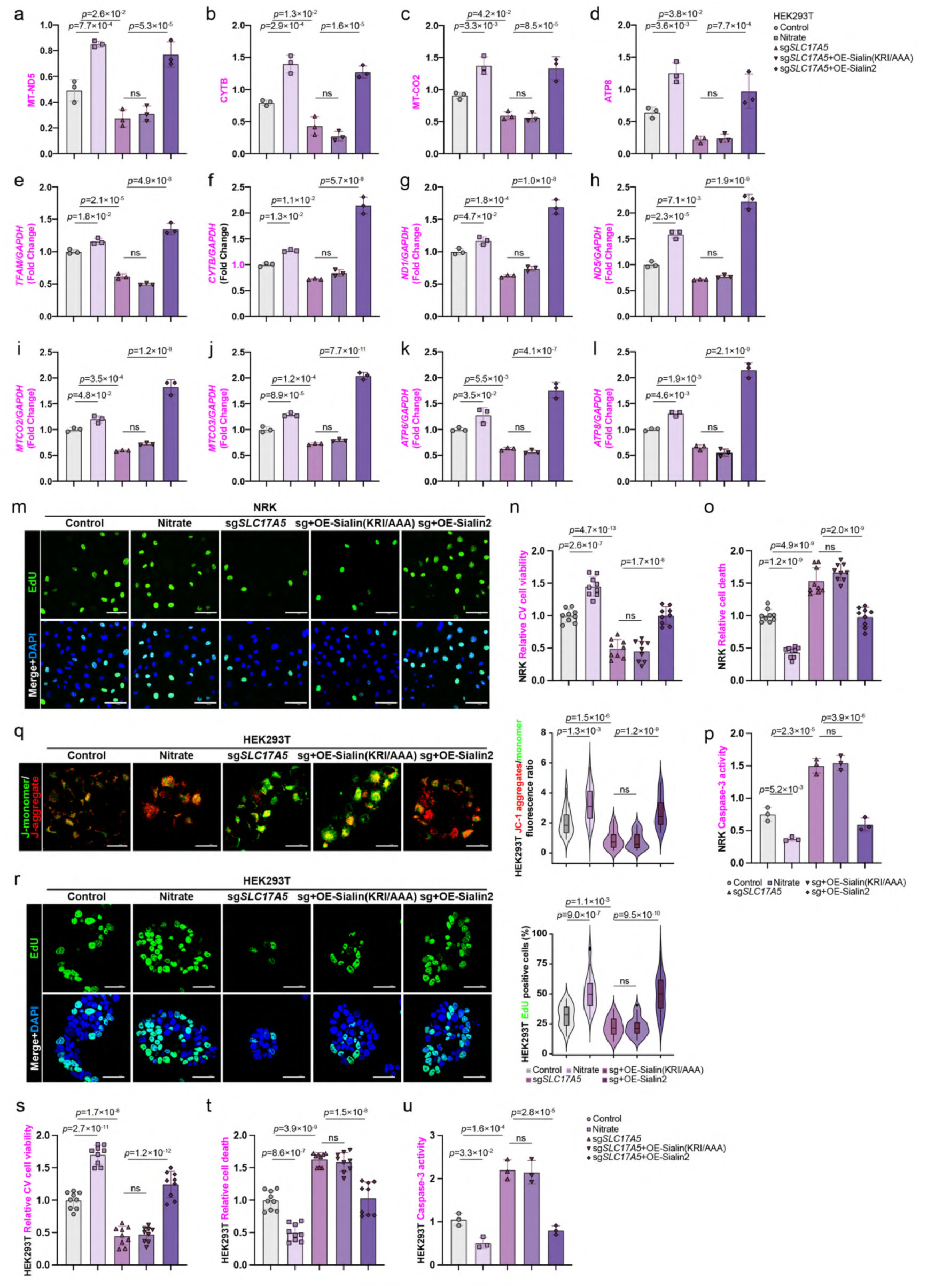
Nitrate-Sialin2 maintains mitochondrial function and cell viability a–d, Immunoblot analysis of mitochondrial-encoded proteins (MT-ND5, CYTB, MT- CO2, and ATP8) in control and sg*SLC17A5* HEK293T cells reconstituted with Sialin (KRI/AAA) or Sialin2 and treated with nitrate (4 mM, 4 h). See immunoblot images in Fig. 4a. e–l, RT-qPCR of mitochondrial genes (*TFAM*, *CYTB*, *ND1*, *ND5*, *MTCO2*, *MTCO3*, *ATP6*, and *ATP8*) in control and sg*SLC17A5* HEK293T cells reconstituted with Sialin (KRI/AAA) or Sialin2 and treated with nitrate (4 mM, 4 h). N = 3. m, IF staining images of EdU in control and sg*SLC17A5* NRK cells reconstituted with Sialin (KRI/AAA) or Sialin2 and treated with nitrate (4 mM, 4 h). N = 30 cells from representative experiments of three repeats. See quantitation in Fig. 4g. n, o, Crystal violet (CV) viability assay (n) and cell death detection (o) in control and sg*SLC17A5* NRK cells reconstituted with Sialin (KRI/AAA) or Sialin2 and treated with nitrate (4 mM, 4 h). Date normalized to control. N = 3 from three independent experiments, each in triplicate. p, Caspase-3 activity assay in control and sg*SLC17A5* NRK cells reconstituted with Sialin (KRI/AAA) or Sialin2 and treated with nitrate (4 mM, 4 h). Activity normalized to control. N = 3. q, r, IF staining images (left) and quantification (right) of JC-1 (q) and EdU (r) in control and sg*SLC17A5* HEK293T cells reconstituted with Sialin (KRI/AAA) or Sialin2 and treated with nitrate (4 mM, 4 h). N = 30 cells from representative experiments of three repeats. s, t, Crystal violet (CV) viability assay (s) and cell death detection (t) in control and sg*SLC17A5* HEK293T cells reconstituted with Sialin (KRI/AAA) or Sialin2 and treated with nitrate (4 mM, 4 h). Date normalized to control. N = 3 from three independent experiments, each in triplicate. u, Caspase-3 activity assay in control and sg*SLC17A5* HEK293T cells reconstituted with Sialin (KRI/AAA) or Sialin2 and treated with nitrate (4 mM, 4 h). Activity normalized to control. N = 3. For all panels, data are represented as mean ± SD, *P* value denotes *t*-test. Scale bar: 10 μm.

**Figure S13.**
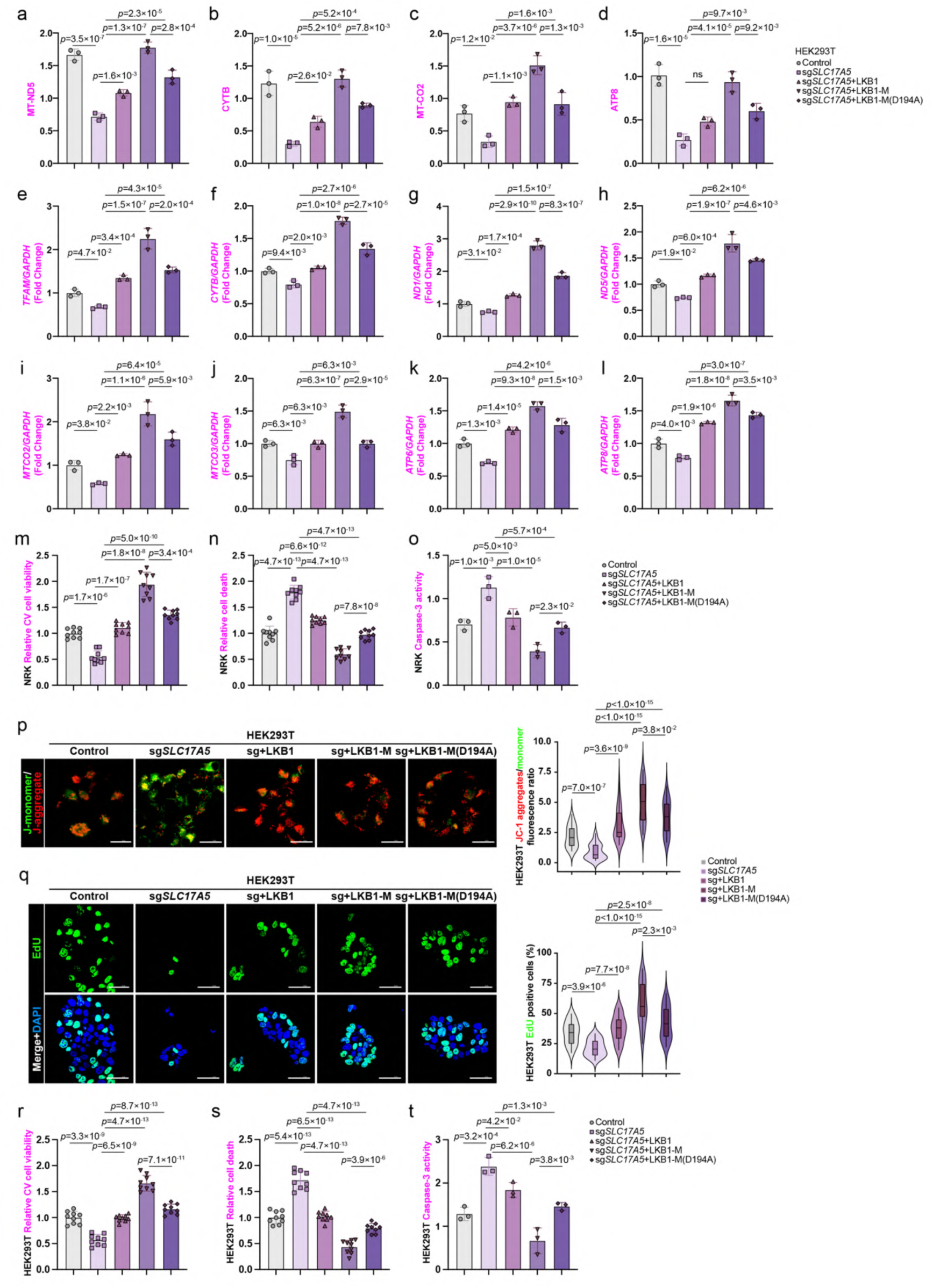
LKB1 mitochondrial recruitment is essential for Sialin2-mediated mitochondrial function and cell viability a–d, Immunoblot analysis of mitochondrial-encoded proteins (MT-ND5, CYTB, MT- CO2, and ATP8) in control and sg*SLC17A5* HEK293T cells reconstituted with vector, Flag-LKB1, Flag-LKB1-M, or Flag-LKB1-M (D194A). See immunoblot images in Fig. 4h. e–l, RT-qPCR of mitochondrial genes (*TFAM*, *CYTB*, *ND1*, *ND5*, *MTCO2*, *MTCO3*, *ATP6*, and *ATP8*) in control and sg*SLC17A5* HEK293T cells reconstituted with vector, Flag-LKB1, Flag-LKB1-M, or Flag-LKB1-M (D194A). N = 3. m, n, Crystal violet (CV) viability assay (m) and cell death detection (n) in control and sg*SLC17A5* NRK cells reconstituted with vector, Flag-LKB1, Flag-LKB1-M, or Flag- LKB1-M (D194A). Date normalized to control. N = 3 from three independent experiments, each in triplicate. o, Caspase-3 activity assay in control and sg*SLC17A5* NRK cells reconstituted with vector, Flag-LKB1, Flag-LKB1-M, or Flag-LKB1-M (D194A). Activity normalized to control. N = 3. p, q, IF staining images (left) and quantification (right) of JC-1 (p) and EdU (q) in control and sg*SLC17A5* HEK293T cells reconstituted with vector, Flag-LKB1, Flag- LKB1-M, or Flag-LKB1-M (D194A). N = 30 cells from representative experiments of three repeats. r, s, Crystal violet (CV) viability assay (r) and cell death detection (s) in control and sg*SLC17A5* HEK293T cells reconstituted with vector, Flag-LKB1, Flag-LKB1-M, or Flag-LKB1-M (D194A). Date normalized to control. N = 3 from three independent experiments, each in triplicate. t, Caspase-3 activity assay in control and sg*SLC17A5* HEK293T cells reconstituted with vector, Flag-LKB1, Flag-LKB1-M, or Flag-LKB1-M (D194A). Activity normalized to control. N = 3. For all panels, data are represented as mean ± SD, *P* value denotes *t*-test. Scale bar: 10 μm.

**Figure S14.**
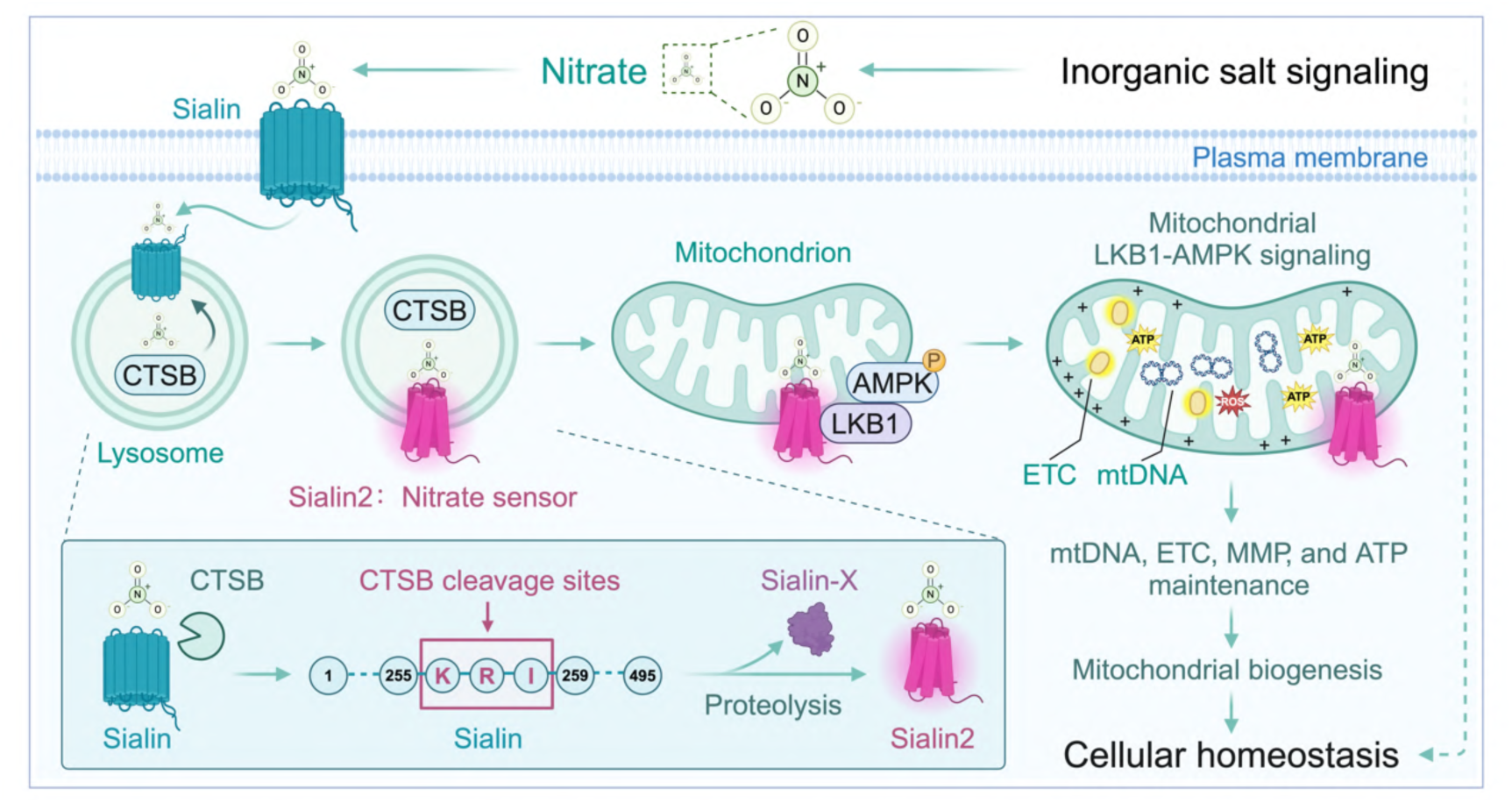
**Sialin2 as the mammalian nitrate sensor coordinates mitochondrial and cellular homeostasis** Summary illustration of the discovery of Sialin2-mediated nitrate sensing and its role in cellular homeostasis. Nitrate triggers CTSB-mediated proteolytic cleavage of the nitrate transporter Sialin, generating two fragments: (i) Sialin2, a mitochondria- localized high-affinity nitrate sensor, and (ii) an uncharacterized fragment (Sialin-X). Sialin2 recruits LKB1-AMPK signaling to mitochondrial microdomains, directly coupling nitrate sensing to mitochondrial homeostasis. This mechanism establishes nitrate as a direct signaling molecule that integrates extracellular nutrient sensing with mitochondrial regulation, establishing the framework of “inorganic salt signaling biology” to maintain cellular homeostasis independently of its classical metabolic roles.

**Table S1.**
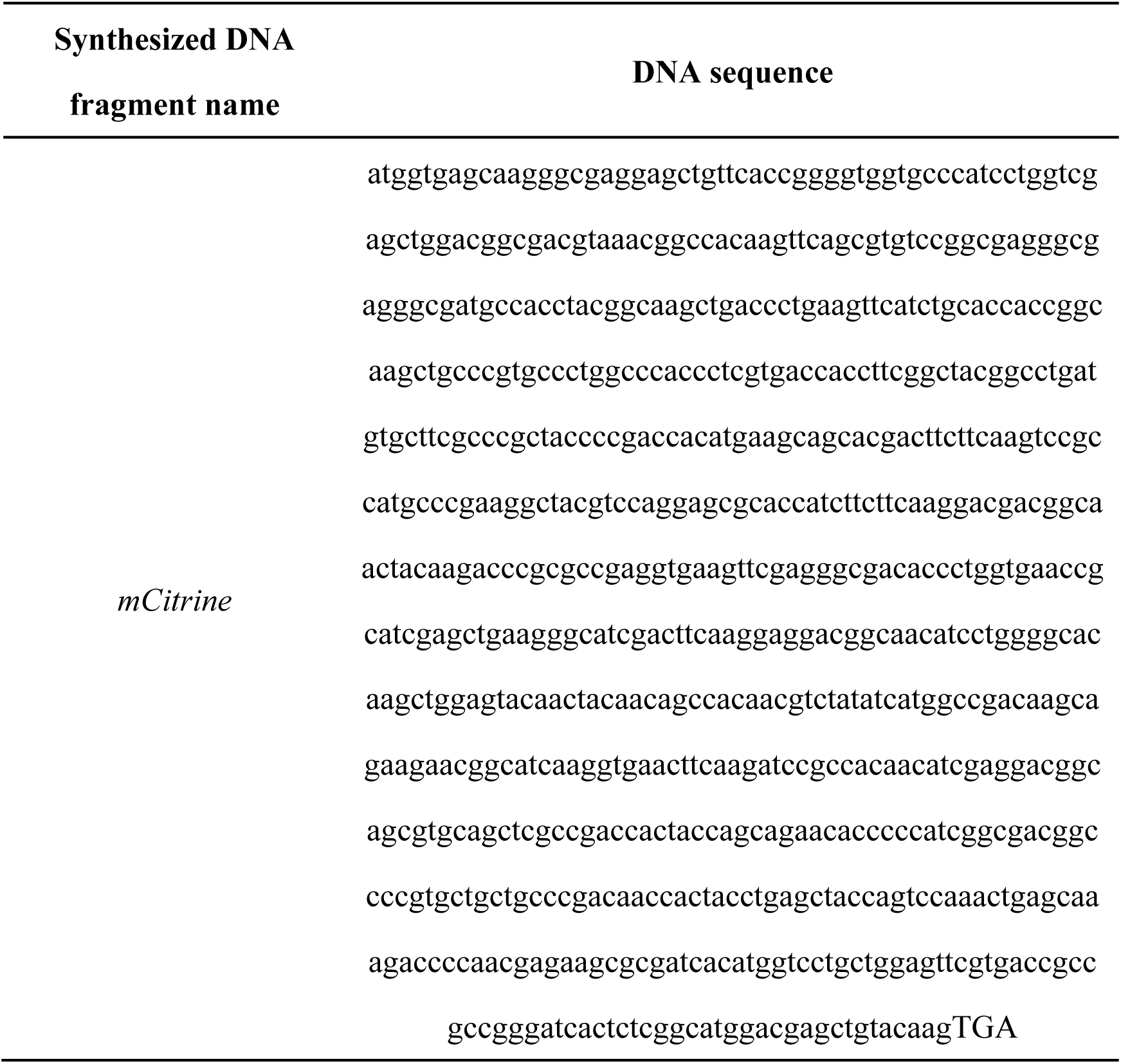
The synthesized DNA sequence.

**Table S2.**
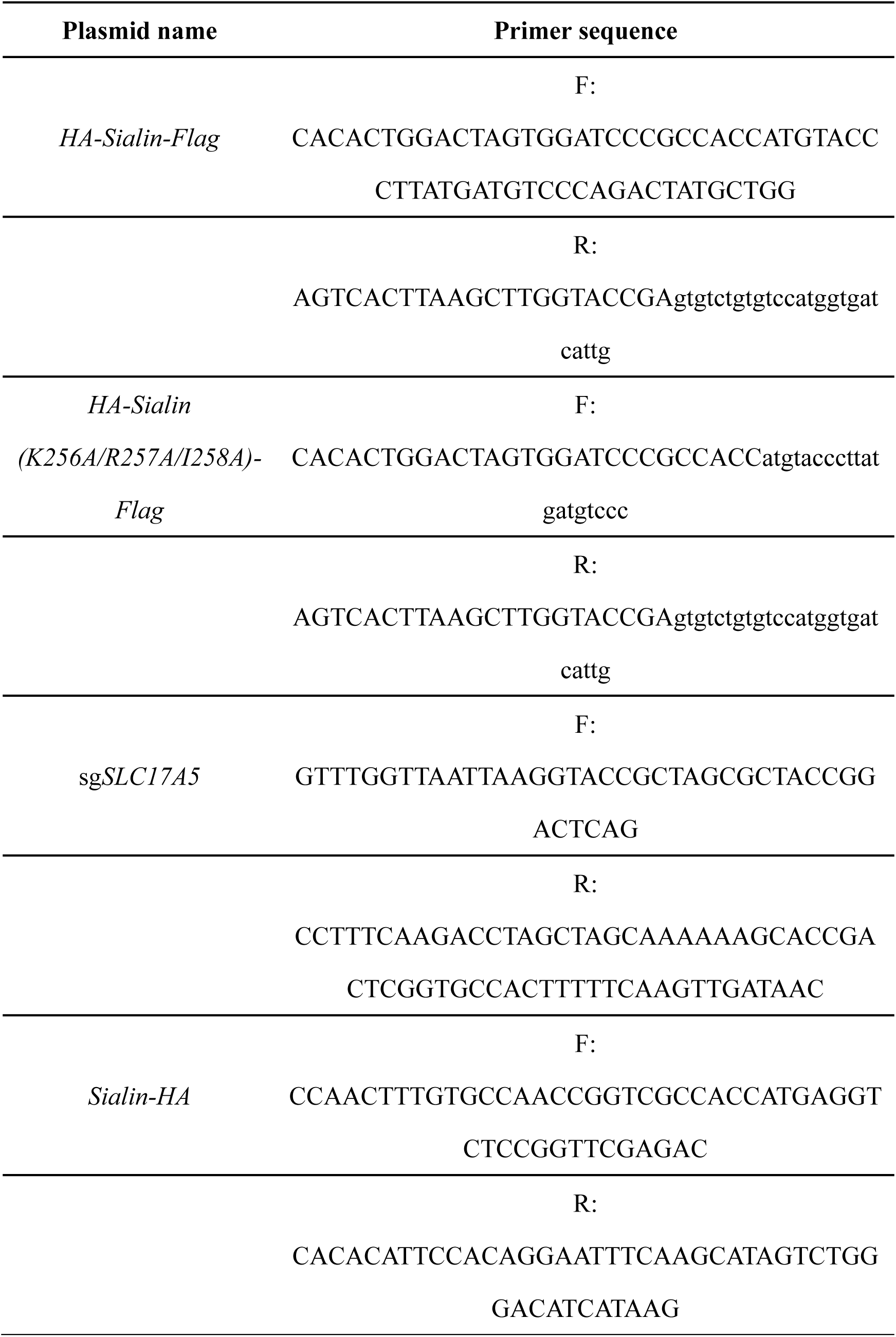
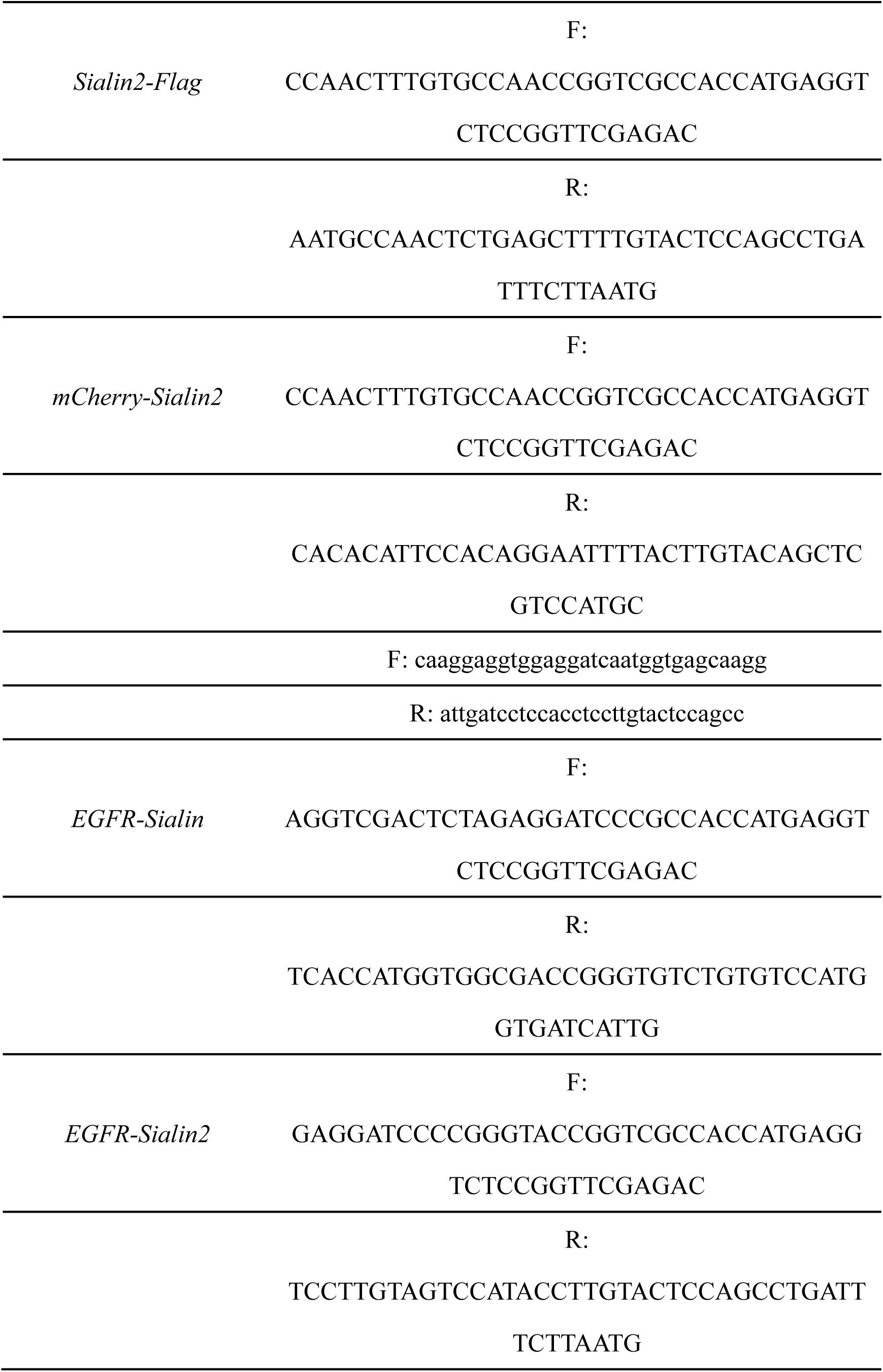
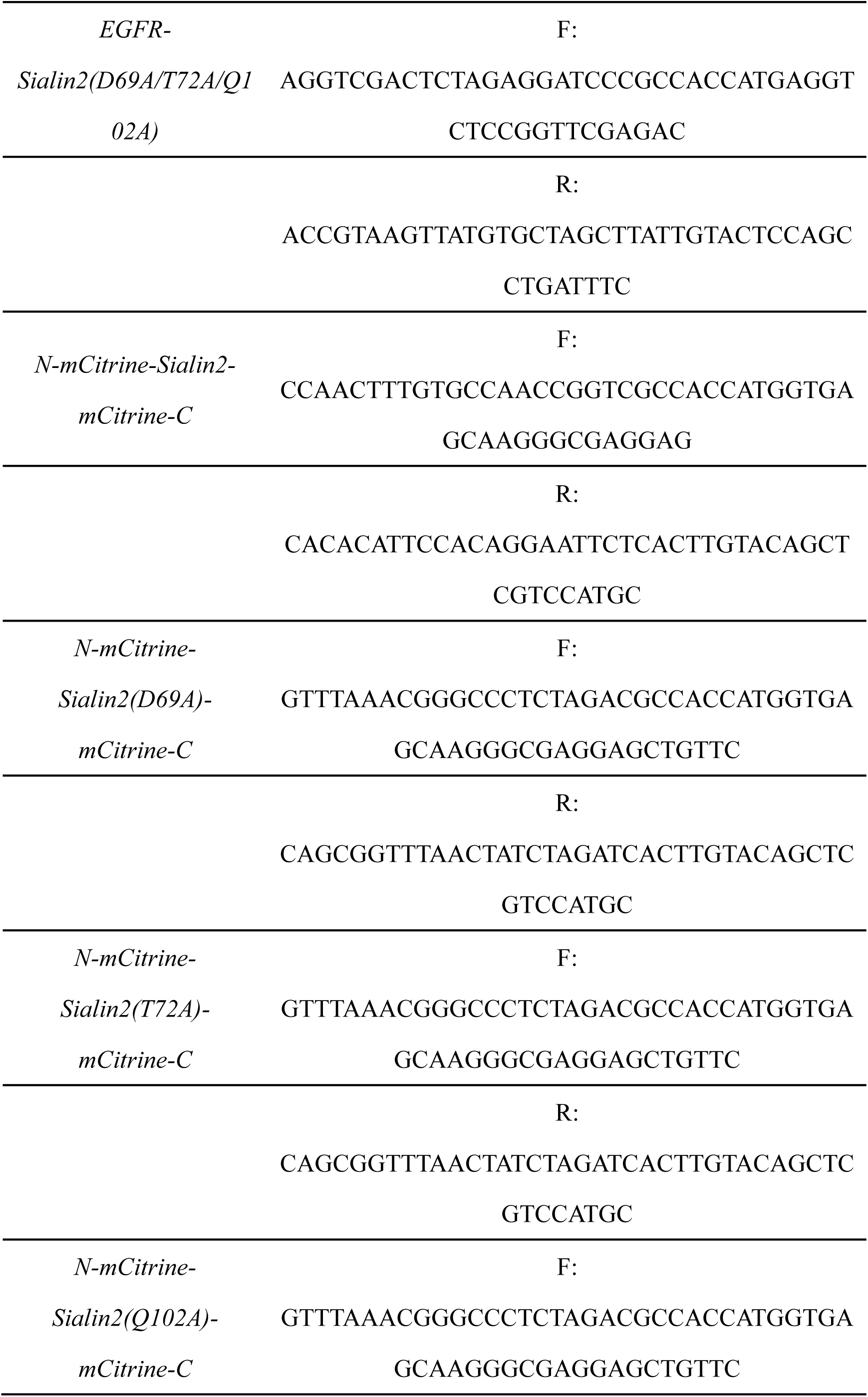
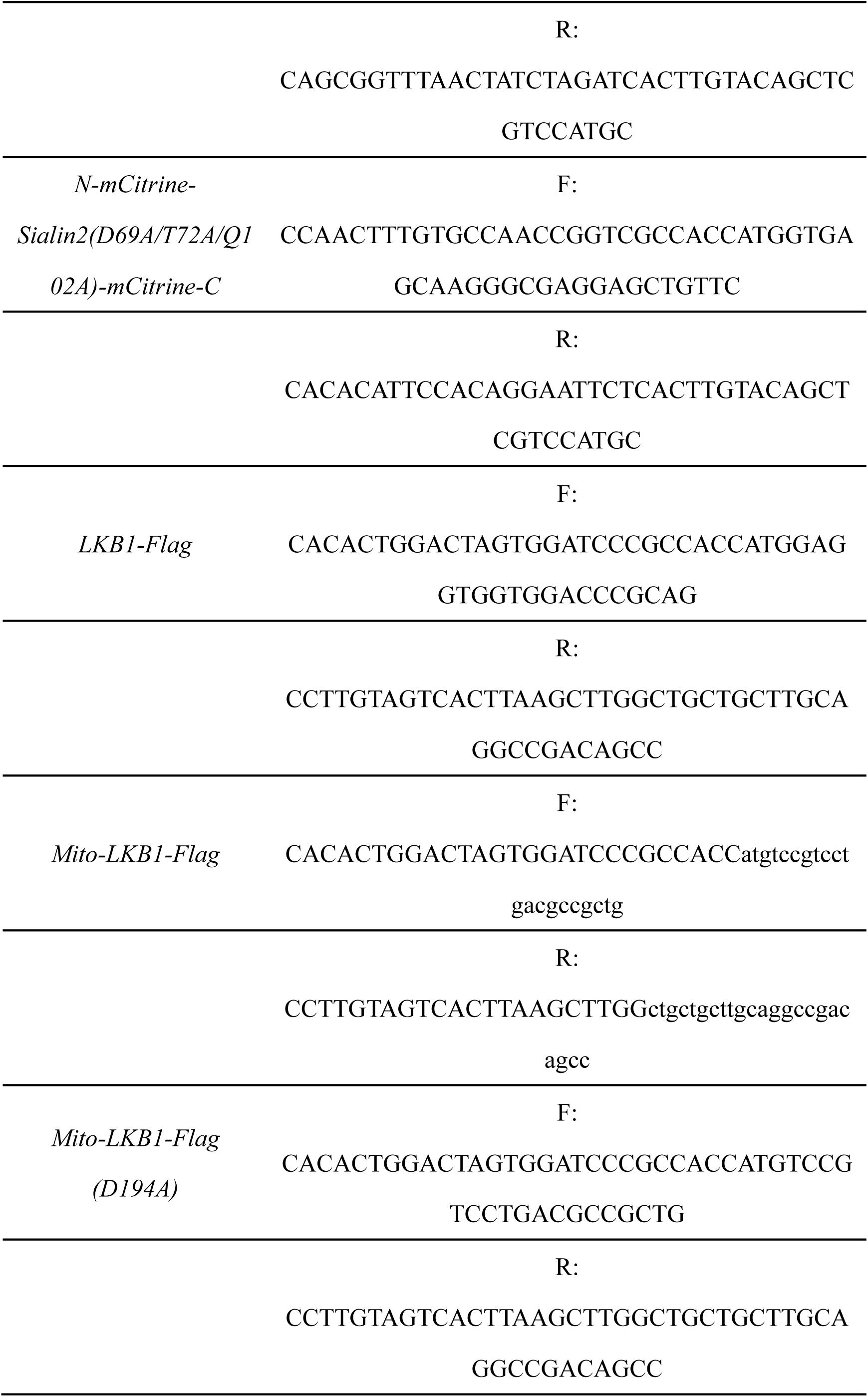
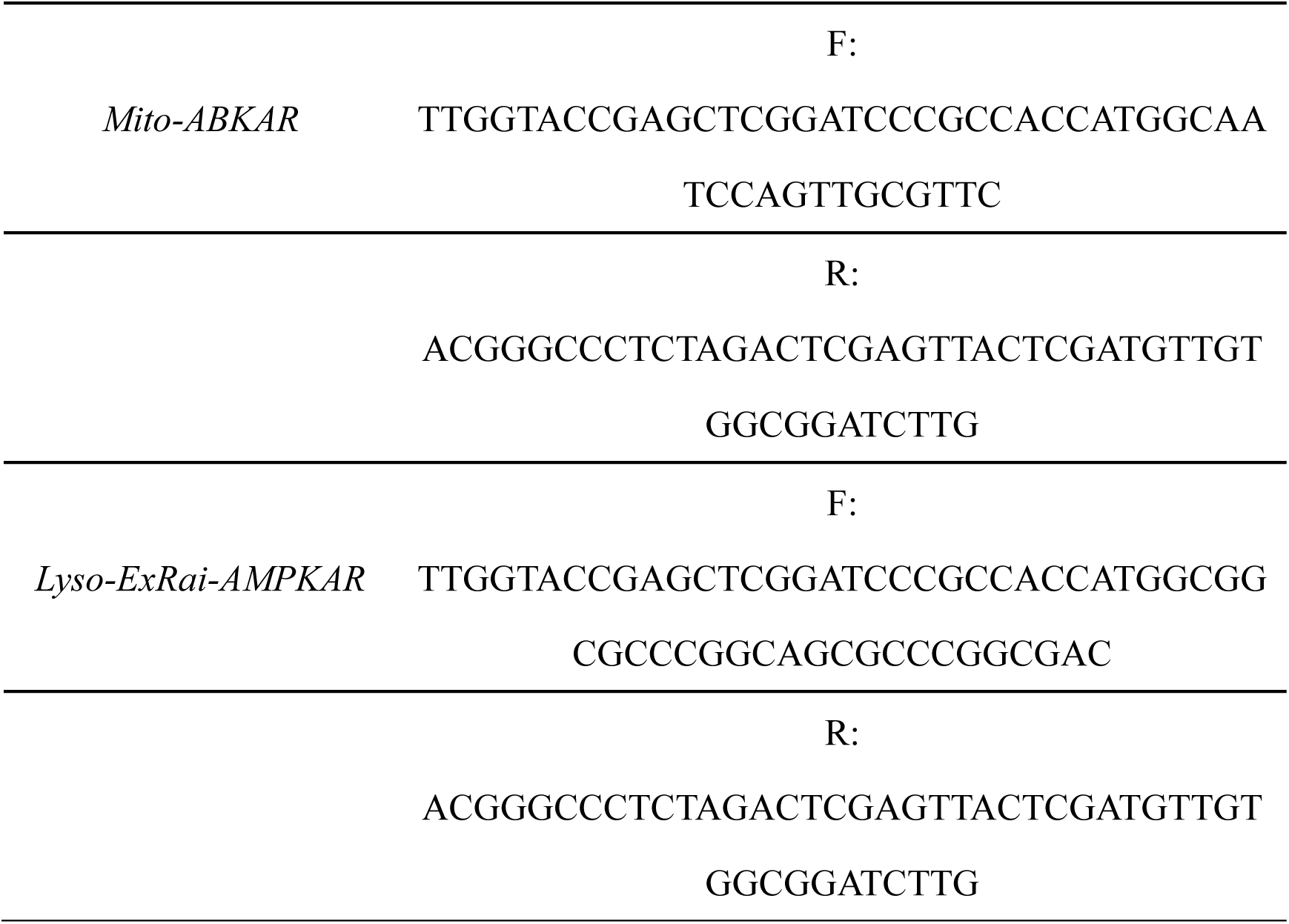
The synthesized DNA plasmid sequences.

**Table S3.**
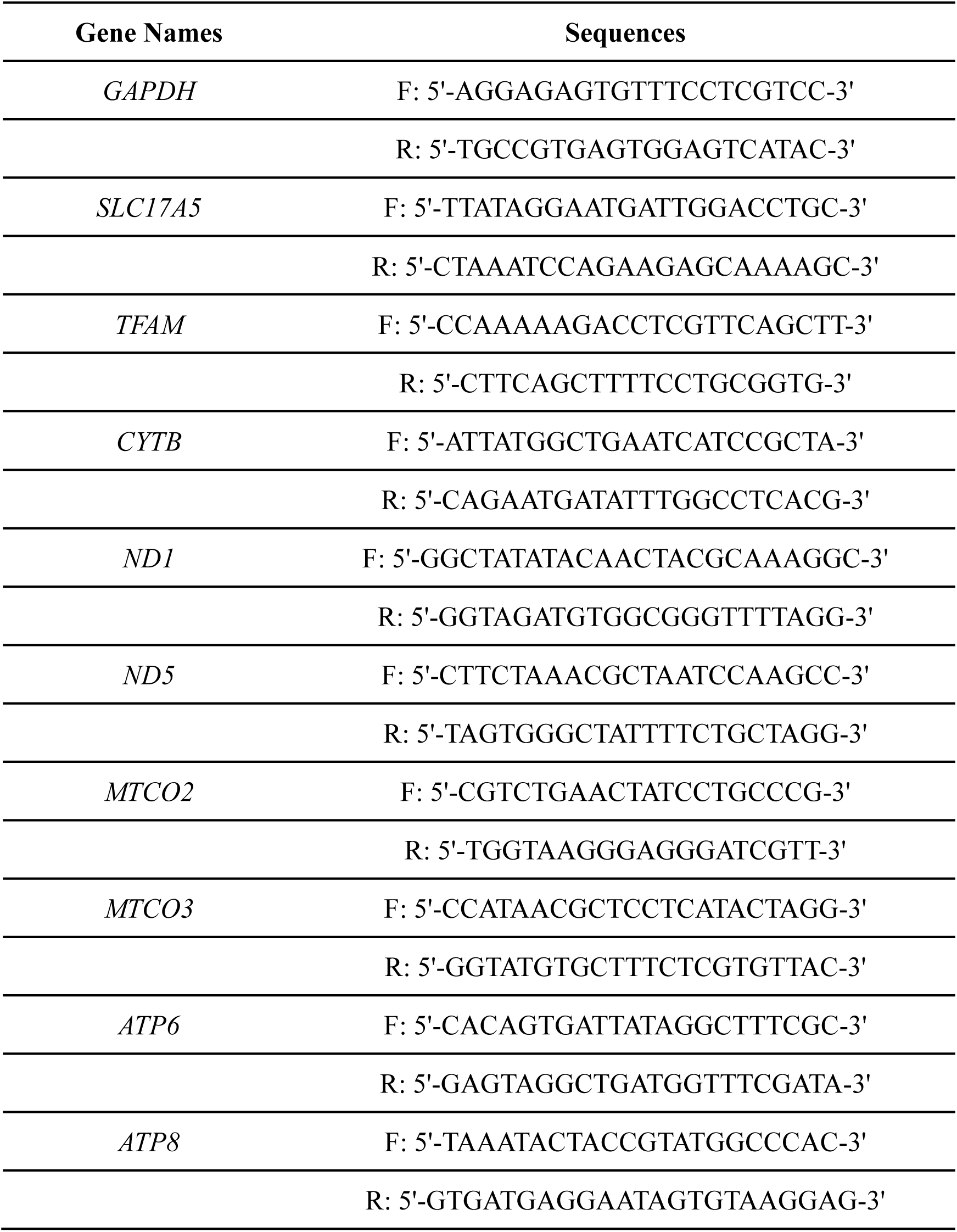
Gene primers.

## References

1. Morris S. M., Jr. Regulation of enzymes of the urea cycle and arginine metabolism. Annual. Rev. Nutr. 22, 87–105 (2002).

2. Lundberg, J. O., Weitzberg, E., & Gladwin, M. T. The nitrate-nitrite-nitric oxide pathway in physiology and therapeutics. Nat. Rev. Drug. Discov. 7(2), 156–167 (2008).

3. Lundberg, J. O., & Weitzberg, E. Nitric oxide signaling in health and disease. Cell 185(16), 2853–2878 (2022).

4. Jin, L., et al. Active secretion and protective effect of salivary nitrate against stress in human volunteers and rats. Free. Radic. Biol. Med. 57, 61–67 (2013).

5. Hu, L., et al. Nitrate ameliorates dextran sodium sulfate-induced colitis by regulating the homeostasis of the intestinal microbiota. Free. Radic. Biol. Med. 152, 609–621 (2020).

6. Ma, L., et al. Rebalancing glucolipid metabolism and gut microbiome dysbiosis by nitrate-dependent alleviation of high-fat diet-induced obesity. BMJ. Open. Diab. Res. Ca. 8, e001255 (2020).

7. Feng, Y., et al. Nitrate increases cisplatin chemosensitivity of oral squamous cell carcinoma via REDD1/AKT signaling pathway. Sci. China. Life. Sci. 64, 1814– 1828 (2021).

8. Feng, X., et al. Dietary nitrate supplementation prevents radiotherapy-induced xerostomia. eLife 10, e70710 (2021).

9. Li, S., et al. Inorganic nitrate alleviates irradiation-induced salivary gland damage by inhibiting pyroptosis. Free. Radic. Biol. Med. 175, 130–140 (2021).

10. Pan, W., et al. Nanonitrator: novel enhancer of inorganic nitrate’s protective effects, predicated on swarm learning approach. Sci. Bull. 68(8), 838–850 (2023).

11. Nilkens, S., et al. Nitrate/oxygen co-sensing by an NreA/NreB sensor complex of Staphylococcus carnosus. Mol. Microbiol. 91, 381–393 (2014).

12. Hu, B., et al. Nitrate–NRT1.1B–SPX4 cascade integrates nitrogen and phosphorus signalling networks in plants. Nat. Plants. 5, 401–413 (2019).

13. Liu, K. H., et al. NIN-like protein 7 transcription factor is a plant nitrate sensor. Science 337, 1419–1425 (2022).

14. Qin, L., et al. Sialin (*SLC17A5*) functions as a nitrate transporter in the plasma membrane. P. Natl. Acad. Sci. USA. 109, 13434–13439 (2012).

15. Martina, J. A., Raben, N., & Puertollano, R. SnapShot: Lysosomal Storage Diseases. Cell 180(3), 602–602.e1 (2022).

16. Schmiege, P., Donnelly, L., Elghobashi-Meinhardt, N., Lee, C. H., & Li, X. Structure and inhibition of the human lysosomal transporter Sialin. Nat. Commun. 15(1), 4386 (2024).

17. Chen, X., et al. Nitrate ameliorates myelin loss and cognitive impairment in Alzheimer’s disease through upregulation of neuronal sialin and subsequent inhibition of TPPP phosphorylation. Sci. Bull. S2095–9273(25)00245-2 (2025).

18. Lundberg, J. O., Carlström, M., & Weitzberg, E. Metabolic effects of dietary nitrate in health and disease. Cell. Metab. 28, 9–22 (2018).

19. de Crom, T. O. E., et al. Dietary nitrate intake in relation to the risk of dementia and imaging markers of vascular brain health: a population-based study. Am. J. Clin. Nutr. 118, 352–359 (2023).

20. Li, X., et al. Salivary nitrate prevents osteoporosis via regulating bone marrow mesenchymal stem cells proliferation and differentiation. J. Orthop. Translat. 45, 188–196 (2024).

21. Dikic I. Proteasomal and Autophagic Degradation Systems. Annu. Rev. Biochem. 86, 193–224 (2017).

22. Olson, O. C., & Joyce, J. A. Cysteine cathepsin proteases: regulators of cancer progression and therapeutic response. Nat. Rev. Cancer. 15(12), 712–729 (2015).

23. Mort, J. S., & Buttle, D. J. Cathepsin B. Int. J. Biochem. Cell. Biol. 29(5), 715–720 (1997).

24. Chacinska, A., Koehler, C. M., Milenkovic, D., Lithgow, T., & Pfanner, N. Importing mitochondrial proteins: machineries and mechanisms. Cell 138(4), 628– 644 (2009).

25. Wong, Y. C., Ysselstein, D., & Krainc, D. Mitochondria-lysosome contacts regulate mitochondrial fission via RAB7 GTP hydrolysis. Nature 554(7692), 382– 386 (2018).

26. Niu, Y., et al. Structural mechanism of SGLT1 inhibitors. Nat. Commun. 13(1), 6440 (2022).

27. Wienken, C. J., Baaske, P., Rothbauer, U., Braun, D., & Duhr, S. Protein-binding assays in biological liquids using microscale thermophoresis. Nat. Commun. 1, 100 (2010).

28. Parker, J. L., & Newstead, S. Molecular basis of nitrate uptake by the plant nitrate transporter NRT1.1. Nature 507, 68–72 (2014).

29. Cabantous, S., et al. A new protein-protein interaction sensor based on tripartite split-GFP association. Sci. Rep. 3, 2854 (2013).

30. Jumper, J., et al. Highly accurate protein structure prediction with AlphaFold. Nature 596(7873), 583–589 (2021).

31. Chatterjee, A., et al. Improving the generalizability of protein-ligand binding predictions with AI-Bind. Nat. Commun. 14, 1989 (2023).

32. Bouguyon, E., Gojon, A., & Nacry, P. Nitrate sensing and signaling in plants. Semin. Cell. Dev. Biol. 23(6), 648–654 (2012).

33. Hardie, D. G., Ross, F. A., & Hawley, S. A. AMPK: a nutrient and energy sensor that maintains energy homeostasis. Nat. Rev. Mol. Cell. Biol. 13(4), 251–262 (2012).

34. Herzig, S., & Shaw, R. J. AMPK: guardian of metabolism and mitochondrial homeostasis. Nat. Rev. Mol. Cell. Biol. 19(2), 121–135 (2018).

35. Miyamoto, T., et al. Compartmentalized AMPK signaling illuminated by genetically encoded molecular sensors and actuators. Cell. Rep. 11(4), 657–670 (2015).

36. Ma, T., et al. Low-dose metformin targets the lysosomal AMPK pathway through PEN2. Nature 603(7899), 159–165 (2022).

37. Toyama, E. Q., et al. Metabolism. AMP-activated protein kinase mediates mitochondrial fission in response to energy stress. Science 351(6270), 275–281 (2016).

38. Shackelford, D. B., & Shaw, R. J. The LKB1-AMPK pathway: metabolism and growth control in tumour suppression. Nat. Rev. Cancer. 9(8), 563–575 (2009).

39. Bazhin, A. A., et al. A bioluminescent probe for longitudinal monitoring of mitochondrial membrane potential. Nat. Chem. Biol. 16(12), 1385–1393 (2020).

40. Shackelford, D. B., et al. LKB1 inactivation dictates therapeutic response of non- small cell lung cancer to the metabolism drug phenformin. Cancer. Cell. 23(2), 143–158 (2013).

41. Malik, N., et al. Induction of lysosomal and mitochondrial biogenesis by AMPK phosphorylation of FNIP1. Science 380(6642), eabj5559 (2023).

42. Jäger, S., Handschin, C., St-Pierre, J., & Spiegelman, B. M. AMP-activated protein kinase (AMPK) action in skeletal muscle via direct phosphorylation of PGC- 1alpha. P. Natl. Acad. Sci. USA. 104(29), 12017–12022 (2007).

43. Bennett, C. F., Latorre-Muro, P., & Puigserver, P. Mechanisms of mitochondrial respiratory adaptation. Nat. Rev. Mol. Cell. Biol. 23(12), 817–835 (2022).

44. Goodson, J. R., et al. An autoinhibitory mechanism controls RNA-binding activity of the nitrate-sensing protein NasR. Mol. Microbiol. 114(2), 348–360 (2020).

45. Hu, B., et al. Nitrate-NRT1.1B-SPX4 cascade integrates nitrogen and phosphorus signalling networks in plants. Nat. Plants. 5(4), 401–413 (2019).

46. Efeyan, A., Comb, W. C., & Sabatini, D. M. Nutrient-sensing mechanisms and pathways. Nature 517(7534), 302–310 (2015).

47. Zhu, X., et al. The nutrient-sensing Rag-GTPase complex in B cells controls humoral immunity via TFEB/TFE3-dependent mitochondrial fitness. Nat. Commun. 15(1), 10163 (2024).

48. Chantranupong, L., Wolfson, R. L., & Sabatini, D. M. Nutrient-sensing mechanisms across evolution. Cell 161(1), 6783 (2015).

49. Myers, R. W., et al. Systemic pan-AMPK activator MK-8722 improves glucose homeostasis but induces cardiac hypertrophy. Science 357, 507–511 (2017).

50. Hord, N. G., Tang, Y., & Bryan, N. S. Food sources of nitrates and nitrites: the physiologic context for potential health benefits. Am. J. Clini. Nutr. 90(1), 1–10 (2009).

51. Jewell, J. L., Russell, R. C., & Guan, K. L. Amino acid signalling upstream of mTOR. Nat. Rev. Mol. Cell. Biol. 14(3), 133–139 (2013).

52. Chintapaludi, S. R., et al. Staging Alzheimer’s disease in the brain and retina of B6.APP/PS1 mice by transcriptional profiling. J. Alzheimers. Dis. 73, 1421–1434 (2020).

53. Huang, X., et al. Fast, long-term, super-resolution imaging with Hessian structured illumination microscopy. Nat. Biotechnol. 36, 451–459 (2018).

54. Zhao, W., et al. Sparse deconvolution improves the resolution of live-cell super- resolution fluorescence microscopy. Nat. Biotechnol. 40, 606–617 (2022).

55. Zheng, S. Q., et al. MotionCor2: anisotropic correction of beam-induced motion for improved cryo-electron microscopy. Nat. Methods. 14, 331–332 (2017).

56. Grant, T. & Grigorieff, N. Measuring the optimal exposure for single particle cryo- EM using a 2.6 A reconstruction of rotavirus VP6. eLife 4, e06980 (2015).

57. Zhang, K. Gctf: Real-time CTF determination and correction. J. Struct. Biol. 193, 1–12 (2016).

58. Zivanov, J., et al. New tools for automated high-resolution cryo-EM structure determination in RELION-3. eLife 7, e42166 (2018).

59. Kimanius, D., Forsberg, B.O., Scheres, S.H., & Lindahl, E. Accelerated cryo-EM structure determination with parallelisation using GPUs in RELION-2. eLife 5, e18722 (2016).

60. Scheres, S. H. RELION: implementation of a Bayesian approach to cryo-EM structure determination. J. Struct. Biol. 180, 519–530 (2012).

61. Scheres, S. H. A Bayesian view on cryo-EM structure determination. J. Mol. Biol. 415, 406–418 (2012).

62. Punjani, A., Rubinstein, J. L., Fleet, D. J., & Brubaker, M. A. cryoSPARC: algorithms for rapid unsupervised cryo-EM structure determination. Nat. Methods. 14, 290–296 (2017).

63. Jumper, J., et al. Highly accurate protein structure prediction with AlphaFold. Nature 596, 583–589 (2021).

64. Emsley, P., Lohkamp, B., Scott, W. G., & Cowtan, K. Features and development of Coot. Acta. Crystallogr. D. 66, 486–501 (2010).

65. Casarotto, P. C., et al. Antidepressant drugs act by directly binding to TRKB neurotrophin receptors. Cell 184, 1299–1313 (2021).

66. Huang, Z. M., et al. Identification of a cellularly active SIRT6 allosteric activator. Nat. Chem. Biol. 14, 1118–1126 (2018).

67. Mita, M., et al. Green Fluorescent Protein-Based Glucose Indicators Report Glucose Dynamics in Living Cells. Anal. Chem. 91(7), 4821–4830 (2019).

